# Effect of a retinoic acid analogue on BMP-driven pluripotent stem cell chondrogenesis

**DOI:** 10.1101/2023.06.20.545738

**Authors:** Fabrizio E. Mancini, Paul E.A. Humphreys, Steven Woods, Nicola Bates, Sara Cuvertino, Julieta O’Flaherty, Leela Biant, Marco A.N. Domingos, Susan J. Kimber

**Author notes:** Corresponding Author: Prof Susan J. Kimber.

## Abstract

Osteoarthritis is the most common degenerative joint condition, leading to articular cartilage (AC) degradation, chronic pain and immobility. The lack of appropriate therapies that provide tissue restoration combined with the limited lifespan of joint-replacement implants indicate the need for alternative AC regeneration strategies. Differentiation of human pluripotent stem cells (hPSCs) into AC progenitors may provide a long-term regenerative solution but are still limited due to the continued reliance upon growth factors to recapitulate developmental signalling processes. Recently, TTNPB, a small molecule activator of retinoic acid receptors (RARs), has been shown to be sufficient to guide mesodermal specification and early chondrogenesis of hPSCs. Here, we modified our previous differentiation protocol, by supplementing cells with TTNPB and administering BMP2 at specific times to enhance early development. Transcriptomic analyses indicated that activation of RAR signalling significantly upregulated genes related to limb and embryonic skeletal development in the early stages of the protocol and upregulated genes related to AC development in later stages. Chondroprogenitors obtained from RAPID-E could generate cartilaginous pellets that expressed AC-related matrix proteins such as Lubricin, Aggrecan, and Collagen II. This protocol could lay the foundations for cell therapy strategies for osteoarthritis and improve the understanding of AC development in humans.

## Introduction

Osteoarthritis (OA) is a chronic incurable skeletal disease that leads to degradation of the articular cartilage matrix, resulting in joint dysfunction and inflammation. It has been estimated to affect over 300 million people world-wide, with knee OA being the most prevalent (1). Existing therapies rely mainly on pain management strategies or joint replacements, which have a limited lifespan (2). Cell-based therapies, in which chondrogenic cells are transplanted to synthesise cartilaginous matrix *in situ* and repair damaged tissue, have been proposed as a long-term therapeutic approach. Human pluripotent stem cells (hPSCs), including human embryonic stem cells (hESCs) and induced pluripotent stem cells (hiPSCs), may be favourable cell sources for regenerative medicine approaches for OA, due to their indefinite proliferative potential and differentiation capacity (3,4). Several approaches have been developed for the differentiation of hPSCs towards chondrogenic cells, with most relying upon treatment with cytokines and small molecules that mimic step-wise activation of pathways active during chondrocyte development (5–7). One challenge for stem-cell based therapeutics for OA has been the generation of stable tissue that does not undergo hypertrophy and mineralisation. As articular chondrocytes of the long bone joints arise from limb bud mesenchymal cells from the lateral-plate mesoderm, we considered that recapitulation of limb and joint developmental signals are likely to be required to obtain true embryonic articular chondrocyte-like cells (6,8,9).

Bone morphogenetic protein (BMP) signalling is important for mesodermal patterning during early embryogenesis, specifying the lateral plate and driving limb bud outgrowth (10–12). Moreover, temporally and spatially regulated BMP signalling is critical throughout chondrogenesis, driving expression and interacting with key chondrogenic transcription factors. BMP signals stimulate differentiation of proliferating chondrocytes which contribute to both the formation of transient growth plate cartilage and to the interzone, which will form the articular (permanent) cartilage. They regulate transcription leading to the synthesis of extra-cellular matrix (ECM) of the cartilage anlage (13–15). Allowing for the prime importance of BMP signals for terminal hypertrophic differentiation of chondrocytes from murine and other models, various studies have demonstrated that it is also an essential ingredient at certain points for hPSC differentiation protocols aimed at generating articular cartilage (AC) (5,6,16– 18). Our group established a refined directed differentiation protocol (RAPID) that recapitulates the developmental systems that regulate limb bud mesenchymal development (6). Cells derived from the RAPID protocol have demonstrated capacity to generate pre-chondrocyte and pre-osteoblast cells, indicating the presence of an osteochondral progenitor population. Pre-chondrocyte cells could be further cultured to generate cartilaginous pellets that expressed type II collagen, aggrecan and had surface localisation of lubricin. Although the RAPID protocol, and other similar chondrogenic hPSC differentiation protocols, rely upon the addition of BMP growth factors to drive differentiation, we have found BMPs to be subject to high batch-to-batch variability.

A recent study found that the use of two molecules, WNT pathway agonist CHIR99021 and retinoic acid receptor (RAR) pan agonist TTNPB, was sufficient to induce chondrogenic differentiation of hPSCs without the use of variable and costly growth factors (19). The authors suggest that chondrogenic differentiation was primarily driven by TTNPB stimulation, in which active nuclear RAR complexes bound to a variety of enhancers and altered chromatin accessibility to promote mesodermal and chondrogenic differentiation. Although RA signalling has been demonstrated to suppress SOX9 expression and antagonise BMP signals that drive chondrogenesis (20–22), it also controls expression of a variety of homeobox (HOX) genes and other genes crucial in early cell-fate specification, including in the limb (23–25). Kawata and colleagues demonstrated via microarray that native *BMP4* and *BMP5* were significantly upregulated upon CHIR99021 and TTNPB stimulation, which then may have facilitated chondrogenic differentiation via enhanced chromatin accessibility. Thus, we investigated the adjustment of BMP signalling activation in the context of RAR activation during differentiation, hypothesising that increased accessibility of chondrogenic genes may facilitate the efficiency of BMP signalling and result in a more robust and representative model of limb-skeletal development and enhanced chondrogenesis.

## Materials and methods

### Culture and Maintenance of human pluripotent stem cells in feeder/xeno-free conditions

MAN13 hESCs were expanded in 6-well cell culture plates previously coated with 5µg/ml Human Recombinant Vitronectin (Gibco, #A27940) in Dulbecco’s Phosphate Buffered Saline (PBS) modified without calcium chloride and magnesium chloride (Sigma-Aldrich, #D8537). Cells were cultured in mTeSR™1 basal medium (StemCell Technologies, #85851) with 20% O_2_ and 5% CO_2_ at 37°C. Medium was replaced every 48 hours, and cells passaged when 80-90% confluent. Cells were dissociated using 0.5mM Ultrapure™ EDTA (Invitrogen, #15575-638) in PBS at 37°C for 3-5 minutes. HiPSCs were grown in 6-well plates pre-coated with 5µg/ml human recombinant Vitronectin in PBS. The cells were maintained in TeSR™-E8™ medium (E8) (StemCell Technologies, #05990) at 20% O2 and 5% CO2 at 37°C, E8 medium was replaced every other day. Cells were passaged every 4-7 days as described previously for hESCs.

### 2D Chondrogenic Differentiation of hESCs

The RAPID-E protocol was adapted from our previously published protocol (6), in which TTNPB was supplemented daily into growth medium and the timing of BMP2 addition was adjusted. Briefly, cells were passaged onto vitronectin as described previously and cultured until approximately 50-60% confluent. Pluripotency maintenance medium was then removed and replaced with Advanced Differentiation Basal Medium (ADBM), consisting of Advanced DMEM F-12 (Thermo Fisher Scientific, #12634010) supplemented with 1% (v/v) L-Glutamine (Gibco, #25030081), 2% (v/v) B27 Supplement (Gibco, #17504001), 0.1mM β-mercaptoethanol (Gibco, #31350010). Cells were sub-cultured on days 4 and 8 of differentiation onto human fibronectin (Merck Millipore, #FC010) coated plates at a ratio of 1:4. Growth factors and cytokines concentration are reported in supplemental experimental procedures **Supplementary Table 1**.

### 3D Chondrogenic Differentiation of hESCs derived chondroprogenitors

Chondroprogenitor cells at the end of the 2D phase of differentiation (Day 11) were dissociated using TryPLE™ express solution at 37°C for 5 minutes. Meanwhile, 15 ml Falcon Tubes were rinsed with Anti-adherence Rinsing Solution (StemCell Technology, #07010) to prevent cell adhesion to the tube. Cell number was determined using a NucleoCounter NC-200 (Chemometec, #900-0201). Cells were diluted in day 11 medium to achieve 1 x 10^6^ cells/ml, and 1 ml of medium was added to each coated falcon tube. Falcon tubes containing cells were spun at 100xg for 3 minutes and then incubated for 96 hours at 20% O2 and 5% CO2 at 37°C. After incubation, medium was replaced with ADBM containing 50µg/ml L-Ascorbic acid 2-phosphate, 20ng/ml GDF5, 25ng/ml BMP2 and 10ng/ml TGFβ. The medium was refreshed every 72 hours, and pellets were kept in culture for 28 days.

### Histological analysis

Samples were processed using a tissue processor (Leica, #ASP300) and Paraffin-embedded samples were cut into 5μm sections using a microtome (Leica, #RM2255) and mounted on Superfrost Plus Adhesion Microscope Slides (Epredia^TM^, # 12302108). Sections were initially subjected to consecutives washes with xylene and ethanol for deparaffinization and finished by washes with deionised water. Endogenous peroxidase activity was blocked in 1% Hydrogen Peroxide (H_2_O_2_) for 30 minutes. Antigen retrieval was conducted by incubating in Pepsin Reagent solution (Sigma-Aldrich #R2283) at room temperature for 15 minutes. The samples were then blocked for 1 hour with 10% serum of the secondary antibody species at room temperature. The samples were subsequently incubated with primary antibodies (**Supplementary Table 2**), in 10% serum, overnight at 4°C, and then incubated 1 hour with secondary antibodies diluted 1:200 (**Supplementary Table 2**). The secondary antibody signal was amplified by incubation with Vectastain Elite ABC Reagent (Vector labs, PK-6100) for 30 minutes. The signal was developed using SIGMAFAST™ 3,3′-Diaminobenzidine (DAB) (Merck, #D4418) for up to 10 minutes at room temperature. Samples were counterstained incubating for 30 seconds in a Mayer’s Modified Haematoxylin solution (Abcam, #220365).

### Alcian Blue/Eosin staining

Sections deparaffinised as reported above, were placed in Acid-alcohol (1% hydrochloric acid in 70% Ethanol) for 30 seconds and drained briefly. Then they were incubated in 1% Alcian blue for 40 minutes and gently washed in distilled water, until it ran clear. After quick rinses in acid-alcohol and 0.5% ammonium water, sections were placed in 95% ethanol for 1 minutes. Slides were placed in Eosin (Abcam, #246824) for 1 minute and then dehydrated, cleared and mounted.

### Flow cytometry analysis

Cells were dissociated using TryPLE™ at 37°C for 5 minutes and counted as previously described. Cells were then transferred into 1.5 ml Eppendorf tubes at the required concentration (not less than 1.0 x 10^5^ cells per tube). Before proceeding with intracellular staining, cells were first fixed with 4% Paraformaldehyde (Thermo Fisher Scientific, #28908) for 10 minutes on ice, then permeabilised by incubating in 70% methanol at 4°C for 7 minutes. For staining, cells were incubated on ice for 30 minutes in a 5% fetal bovine serum-PBS (FBS-PBS) staining solution containing PRRX1 (Abnova, #H00005396-W01P) primary antibody diluted 1:100. Incubation with Alexa Fluor 488 Goat Anti Rabbit (Thermo Fisher Scientific, #A11008) diluted at 1:400 followed under the same conditions as the primary. After incubation, cells were resuspended in 2% FBS-PBS and analysed by flow cytometry using an LSRFortessa™ (BD) and BD FACSDiva™ Software for Windows (Version 8.0). Cells of interest were identified based on their size and granularity through Forward vs Side Scatter (FSC vs SSC) gating. FSC-Height (FSC-H) vs FSC-Area (FSC-A) plots were used to exclude doublets. Unstained cells and single-stained compensation controls were used to set flow cytometer parameters and gating. At least 10,000 events per condition were collected. Data analysis was performed using FlowJo™ Software for Windows (Version 10.6.8.).

### RNA extraction and gene expression analysis by RT-qPCR

Total RNA was extracted using Monarch® Total RNA Miniprep Kit (New England Biolabs inc. #T2010S) following manufacturer protocol. Reverse-transcription of 2µg of RNA was performed using High Capacity cDNA Reverse Transcription Kit (Thermo Fisher Scientific, #4368814). Gene-specific primers were designed using Primer-BLAST online software (26) and are reported in **Supplementary Table 3**. RT-qPCR for gene expression was assessed using 10 ng of cDNA, gene-specific primers and Power SYBR Green PCR Master Mix (Applied Biosystems, UK, #4309155) with a Bio-Rad C1000Touch™ Thermal Cycler. Gene expression was normalised to *GAPDH,* and relative gene expression was calculated using the 2^-ΔCT^ method. At least three independent biological repeats analysed in triplicate were performed.

### RNA-seq analysis

An analysis was performed on paired-end sequences from an Illumina HiSeq4000 sequencer using FastQC (http://www.bioinformatics.babraham.ac.uk/projects/fastqc/) by the Genomic Technologies Core Facility at the University of Manchester. Adapter sequences were removed and the reads were quality checked using Trimmomatic_0.39 (PMID: 24695404). Reads from the RNA-seq experiment were mapped against the reference human genome (hg38) and gene counts were determined using annotation from GENCODE 39 (http://www.gencodegenes.org) with STAR_2.7.7a (PMID: 23104886). Normalization, Principal Components Analysis, and differential expression were calculated using DESeq2_1.36.0 (PMID: 25516281). Gene enrichment analyses were performed with EnrichR (27–29) submitting significant differentially expressed genes (adjusted p-value<0.05) and upregulated or downregulated by at least 1 fold change (unless specified). Graphs were made using GraphPad Prism 9.4.1 or SRPlot, (http://www.bioinformatics.com.cn/srplot), an online platform for data analysis and visualization. Volcano Plots were made using VolcaNoseR (30). Hierarchical clustering analysis was performed using Gene Cluster software (31).

### Assessment of BMP2 growth factor activity

To assess BMP2 growth factor activity, TC28a2-BRE-PEST cells (previously described (32)) were stimulated with BMP2 at a range of concentrations (0.1-1,000ng/ml). Nano-Glo® live cell assay system (Promega, #N2011) was used for analysis of luminescence after stimulation at different time points. Luminescence was measured in raw luminescent units (RLU) using a GloMax® multimodal plate reader (Promega). Reads across a time course were performed using separate wells.

### Statistical analysis

Statistical analyses were conducted using GraphPad Prism 9.4.1 and data are presented as mean ± standard error of mean (SEM). The selection of statistical test in this study was based on the normality of the data and the number of groups being compared. Normality of the data was evaluated through the Shapiro-Wilk test. Data that met the assumption of normality were analysed using t-test, for two groups, and ANOVA for multiple groups. A p-value of < 0.05 was considered statistically significant.

## Results

### Activation of RAR signalling and specific timing of BMP2 addition improves chondrogenesis

BMP signalling plays a major role in human cartilage development and thus BMP family growth factors comprise a component of many *in vitro* human pluripotent stem cell (hPSC) chondrogenic differentiation protocols (5,6,33). However, high variability of BMP2 activity between batches is a likely cause of unwanted variation in chondrogenic differentiation efficiency (**Supplementary Figure 1**). To further understand the role of BMP and improve chondrogenic differentiation efficiency, we investigated adjusting the timing of BMP addition. Since activation of RAR signalling is crucial for limb development and has previously been shown to efficiently drive chondrogenic differentiation when used after Wnt pathway stimulation (19), we also examined the influence of BMP in the context of RAR activation. Using our established chondrogenic differentiation protocol to drive hPSCs towards limb-derived, pre-articular chondrocytes (6), we firstly evaluated the addition of a selective pan-RAR agonist, TTNPB (4-[(E)-2-(5,6,7,8-tetrahydro-5,5,8,8-tetramethyl-2-naphthalenyl)-1-propenyl]benzoic acid) in the absence of BMP2. Through transcriptional analysis of some key chondrogenic genes, we found that addition of TTNPB throughout the protocol significantly increased expression of *SOX9*, whilst significantly reducing expression of *COL1A1* and *RUNX2* at the end of the protocol (**Figure 1A**). Although expression of *COL2A1* was increased at day 8 compared to pluripotent cells, its expression then fell significantly by day 11.

**Figure 1:**
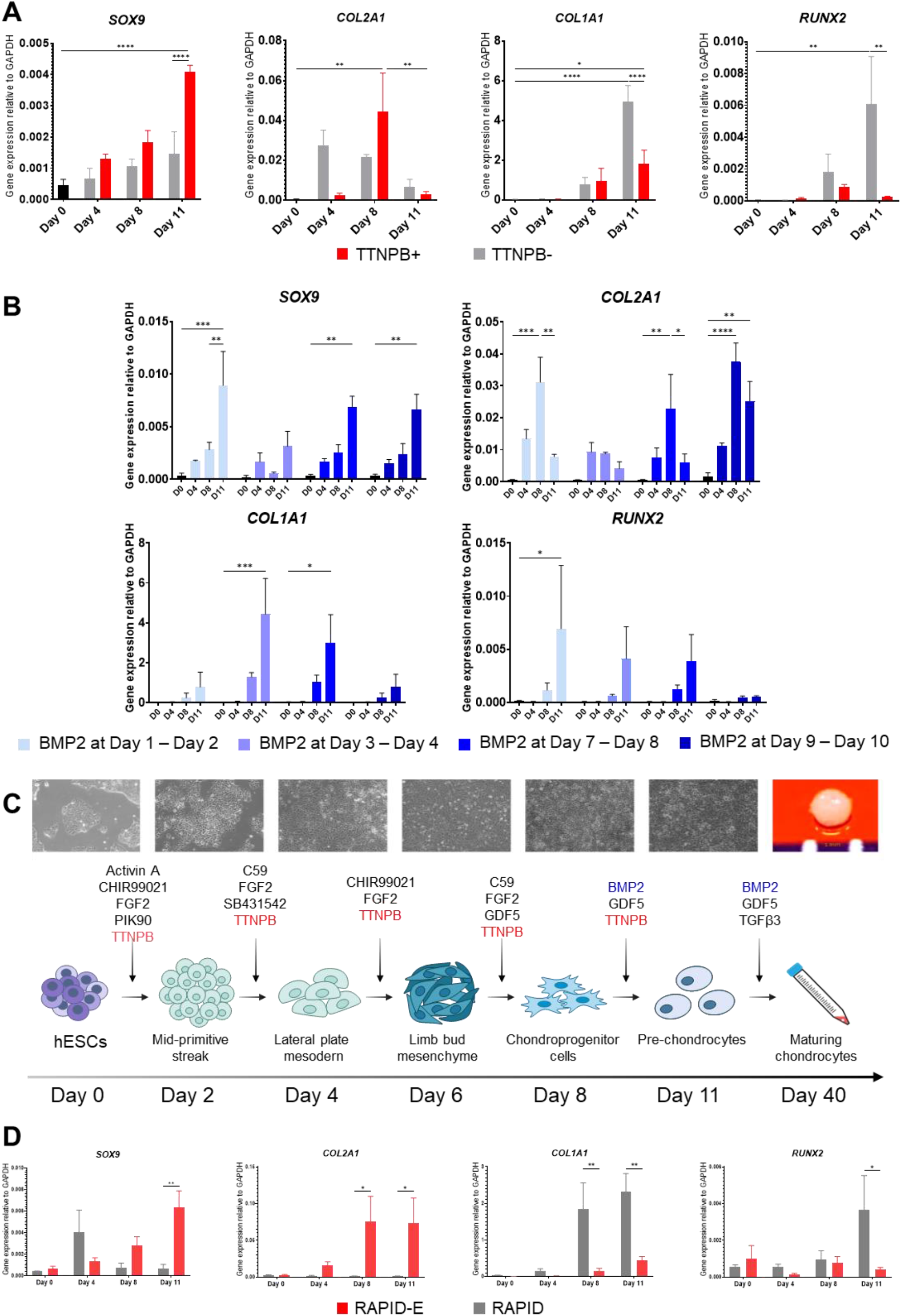
TTNPB supplementation and different BMP2 timing effects on the expression of key chondrogenic genes. **(A)** RT-qPCR analysis of chondrogenic markers expression with and without supplementation of TTNPB and without supplementation of BMP2. Data are presented as the mean value ± SEM of N=4 independent experiments for *SOX9* and N=3 for the other genes. Statistical analysis was performed using an Ordinary two-way ANOVA test (* p<0.05, ** p<0.01, *** p<0.005, **** p < 0.0001). **(B)** RT-qPCR analysis of chondrogenic markers expression for different timing of BMP2 supplementation. Data are presented as the mean value ± SEM of N=3 independent experiments. Statistical analysis was performed using an RM two-way ANOVA test (* p<0.05, ** p<0.01, *** p<0.005, **** p < 0.0001). **(C)** Schematic of the RAPID-E protocol, TTNPB was supplemented during the 2D phase, while BMP2 was supplemented at the last stages of the protocol and throughout the 3D phase. **(D)** RT-qPCR analysis comparing chondrogenic markers expression for hESCs differentiated using the RAPID and the RAPID-E protocols. Data are presented as the mean value ± SEM of N=3 independent experiments. Statistical analysis was performed using a two-way ANOVA (* p<0.05, ** p<0.01). Gene expression levels were normalised to the housekeeping gene *GAPDH*.

As *COL2A1* expression is essential for the generation of cartilaginous ECM and driven by BMP signals (34–36), we investigated adding BMP2 back at different times during differentiation (**Figure 1B**). Addition of BMP2 during the first stage of the protocol (day 1-2) significantly enhanced expression of *SOX9* by day 11, but did not prevent a significant decrease in *COL2A1* expression between day 8 and 11. However, we observed a significant increase in *RUNX2* expression at day 11 compared with no addition of BMP2. Similarly, addition of BMP2 during either day 3-4 or day 7-8 increased expression of *SOX9* and *RUNX2* at day 11, but not *COL2A1*. We also observed an increase in *COL1A1* expression at day 11 with early additions of BMP2. However, when BMP2 was added in the final stages of 2D differentiation (day 9-11), *SOX9* expression was significantly increased, *COL2A1* expression did not fall significantly between day 8 and 11, and we retained relatively low expression of *COL1A1* and *RUNX2* (**Figure 1B**). We selected the protocol with TTNPB from day 1 to 10 and BMP2 from day 9 to 11 for further investigation, this protocol is referred to as RAPID-E from here onwards (**Figure 1C**). When compared to the published RAPID protocol (6), cells differentiated through RAPID-E expressed significantly higher *SOX9* and *COL2A1* combined with significantly lower *COL1A1* and *RUNX2* at day 11 (**Figure 1D**). Cells were sub-cultured twice, which reflected a high proliferation rate up to day 8, with proliferation slowing by day 11 (**Supplementary Figure 2**). To confirm reproducibility of the protocol, we subjected five hiPSC lines from different genetic backgrounds to the modified differentiation protocol (**Supplementary Figure 3**). Gene expression patterns were similar across all cell lines, demonstrating the robustness of differentiation towards chondrogenic cells.

### RNAseq confirms chondrogenic differentiation of pluripotent stem cells with addition of TTNPB and early timed BMP addition

To investigate and characterise lineage specification and chondrogenic ability of our modified differentiation protocol involving TTNPB and BMP2 delivery at optimal stages, we performed RNAseq analysis of hPSCs at day 6 and day 11 with and without TTNPB (**Figure 2A**). Principal component analysis (PCA) indicated distinct separation of the two differentiation stages and treatments, with differentiated cells separated from pluripotent cells on PC1, and samples treated with TTNPB separated from non-treated on PC2 (**Figure 2B**). Hierarchical clustering analysis additionally highlighted the separation of cells treated with TTNPB from non-treated, although cells still clustered based upon differentiation stage (**Supplementary Figure 4**). Differential gene expression (DEG) analysis indicated that D11+T cells significantly upregulated genes critical for skeletal morphogenesis and chondrogenesis, such as *SOX5*, *SOX6*, *SOX9* and *COL2A1*, alongside downregulation of pluripotency markers such as *POU5F1*, *NANOG* and *SOX2* (**Figure 2C**). Additionally, gene ontology (GO) analysis of significantly upregulated DEGs in D11+TTNPB (D11+T) cells in comparison to pluripotent cells enriched for biological process terms associated with chondrogenesis, ECM synthesis and skeletal development (**Figure 2D**). Therefore, the RAPID-E protocol was able to generate a chondrogenic-like population of cells by day 11 of differentiation.

**Figure 2:**
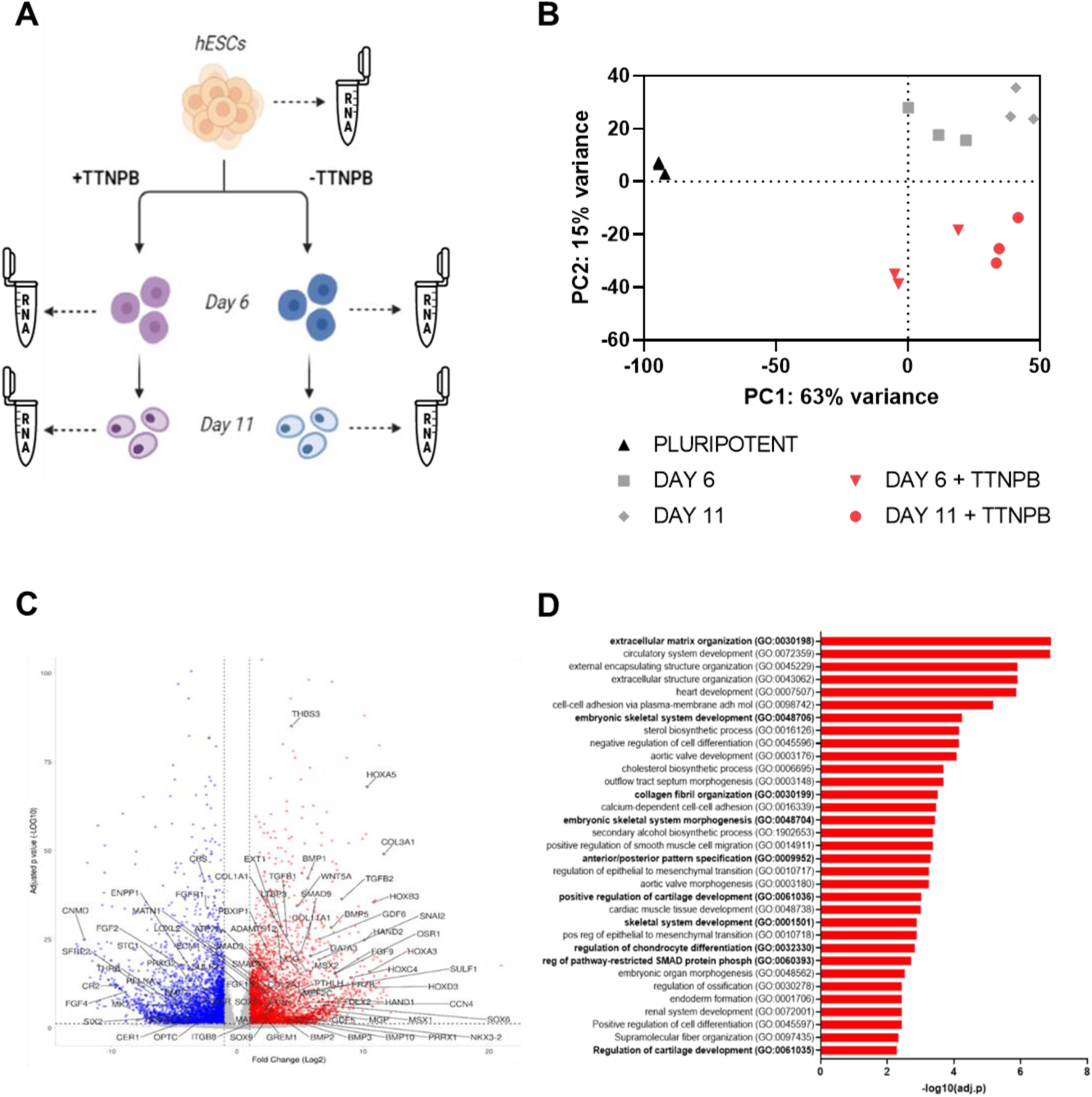
RNAseq analysis of cells differentiated with RAPID-E. **(A)** Cells were differentiated using the RAPID-E with or without addition of TTNPB. Samples were collected at Day 0 (Pluripotency stage), Day 6 (Limb bud induction), and Day 11 (end of chondrogenesis in 2D). **(B)** PCA analysis results showing the distribution of samples on the plane identified by the first (PC1) and the second (PC2) principal components. (**C**) Gene Ontology analysis of biological process enriched in differentially upregulated expressed genes of Day 11 treated samples against day 0 pluripotency samples. Terms in bold are terms related to cartilage and skeletal development. EnrichR was utilised to perform functional enrichment analysis of all upregulated genes with Log2 Fold change > 1 and adjusted p-value < 0.05. **(D)** Volcano plot of differentially expressed genes when comparing differentiated cells at Day 11 with pluripotent cells. Terms in bold are terms related to cartilage development (GO: 0051216). Differentially expressed genes were analysed at a threshold of >1log2 and <-1log2 fold change and an adjusted p-value of >0.1.

### Addition of TTNPB affects lineage specification and chondrogenic capacity

To evaluate the specific effects of TTNPB upon differentiation, we compared cells from our optimised RAPID-E protocol that were treated with TTNPB throughout to those without TTNPB, at day 6 and day 11. Addition of TTNPB induced a large number of transcriptional changes at both stages, with over 1000 significant DEGs (**Supplementary Figure 5A**). To investigate the effect of transcriptional changes induced directly by RAR signalling, we established the commonly up- and down-regulated DEGs when comparing cells treated with or without TTNPB at both day 6 and day 11 (**Figure 3A**). We observed that 259 and 355 shared genes were up- and down-regulated respectively and therefore most likely affected by TTNPB addition. Drug signature database (DSigDB) enrichment analysis of our common upregulated DEGs using EnrichR revealed a significant over representation of gene previously shown to be regulated by Retinoic acid and TTNPB addition, verifying activation of know RA response genes (**Supplementary Figure 5B**). Furthermore, some genes associated with RA signalling were upregulated in samples treated with TTNPB, such as *RARA* and *RARB*, although others were unchanged or diminished, such as *RXRA*, *RXRB*, *RXRG* and *RARG* (**Supplementary Figure 5C**). Importantly however, GO term analysis of upregulated DEGs indicated that addition of TTNPB resulted in enrichment of terms associated with skeletal system morphogenesis, whilst downregulated DEGs enriched for terms associated with cardiac and muscle tissue development (**Figure 3B**). Thus, TTNPB addition appeared to enhance specification of limb bud-like mesenchyme along the skeletal route.

**Figure 3:**
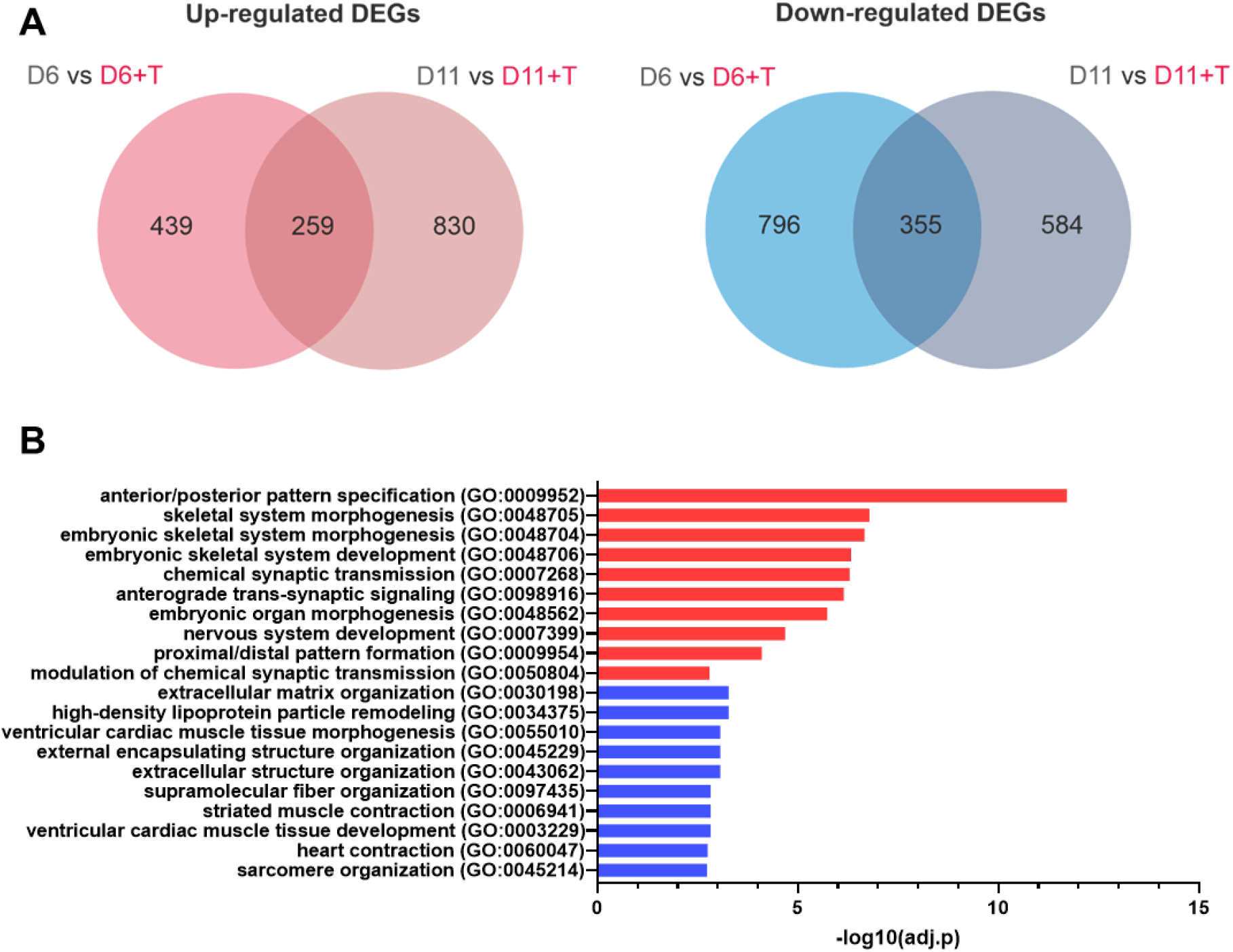
Analysis of transcriptional changes induced by TTNPB addition. (**A**) Venn diagram indicating commonality of genes upregulated (left) or downregulated (right) at Day 6 and 11 when cells were treated with TTNPB. Only genes with >1log2 or <-1log2 fold change and an adjusted p-value of >0.1 were considered. (**B**) Gene Ontology analysis of biological process enriched in differentially commonly expressed genes at day 6 and day 11 of differentiation. EnrichR was utilised to perform functional enrichment analysis.

As cells at day 6 of differentiation should exhibit a limb bud mesenchymal-like phenotype, we investigated differences in lineage marker expression between D6 and D6+T cells (**Figure 4A**). Analysis of lineage markers associated with limb formation indicated that the addition of TTNPB quickened lineage progression, with D6 cells in the absence of TTNPB exhibiting a more premature lateral-plate mesodermal phenotype while cells exposed to TTNPB expressed more genes associated with limb bud mesenchyme (**Figure 4C**). In the presence of TTNPB, more than 70% of the population of D6 cells showed staining for the limb bud mesenchyme marker *PRRX1* (**Supplementary Figure 6).** The addition of TTNPB also appeared to induce significant differences in HOX gene expression by both day 6 and day 11 (**Supplementary Figure 7**). However, most of these HOX genes were associated with anterior embryonic development, such as *HOXA1-5* and *HOXB2-9*, suggesting these clusters were particularly sensitive to retinoic acid (RA) pathway stimulation. Similarly, we analysed differences in expression of individual chondrogenic genes at D11 between cells differentiated with or without TTNPB to confirm the pro-chondrogenic effect of TTNPB. Cells treated with TTNPB exhibited significantly higher expression of *SOX9*, *COL11A1* and *PTHLH*, whilst exhibiting significantly lower expression of *FGFR3*, *RUNX2* and *PTH1R,* genes that are associated with pre-hypertrophic chondrocytes within the growth plate (**Figure 4D**, **Supplementary Figure 8**). Therefore, cells appeared to exhibit a phenotype more associated with resting-zone chondrocytes, suggesting TTNPB helped drive a chondrogenic phenotype.

**Figure 4:**
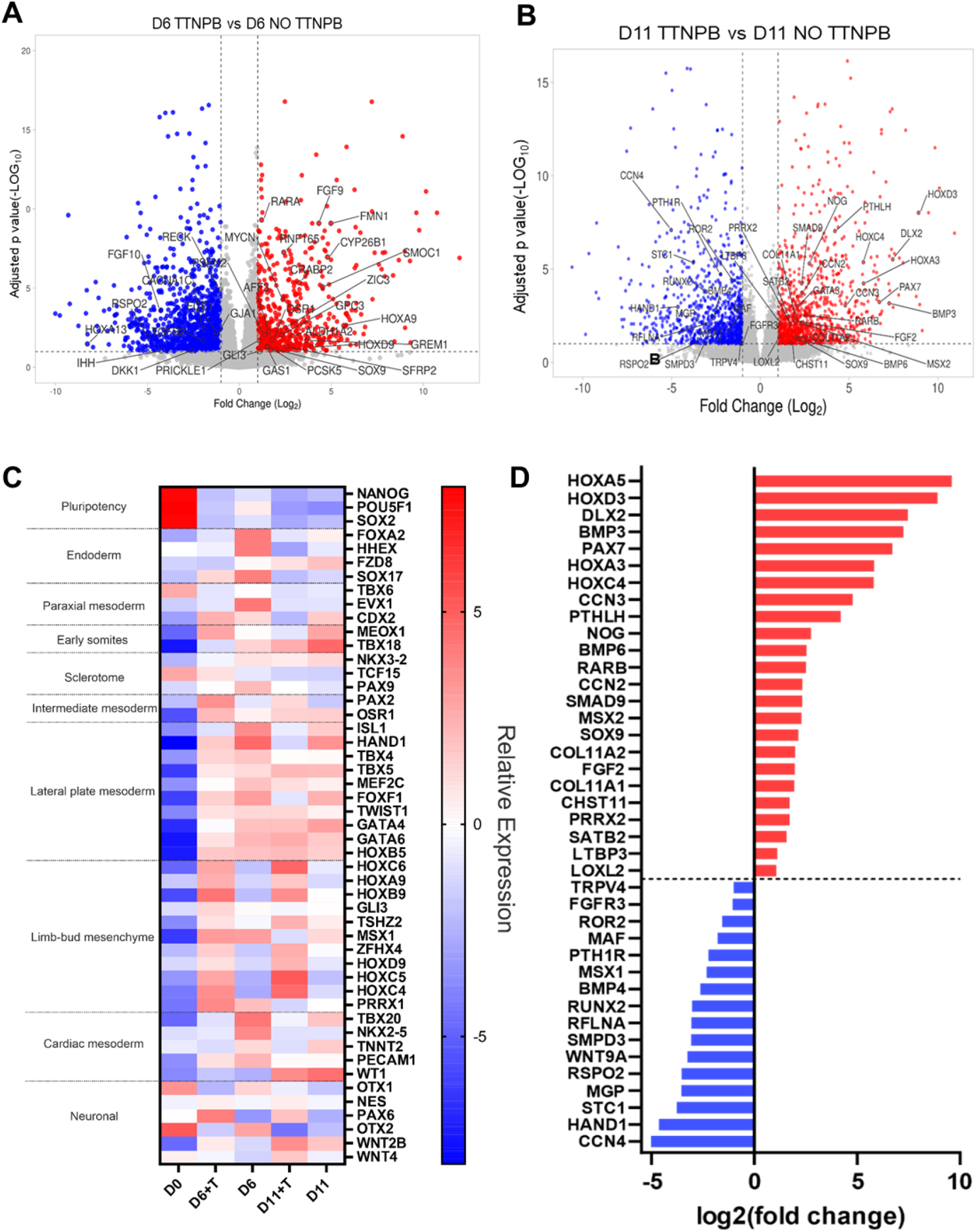
Comparison of differentially expressed genes in RAPID-E differentiated cells either with or without TTNPB. **(A)** Volcano plot analysis of differentially expressed genes in day 6 samples. Red dots are upregulated genes, and blue dots are downregulated. Labelled genes are genes in common with the Limb Development term (GO: 0060173). Differentially expressed genes were analysed at a threshold of >1log2 and <-1log2 fold change and an adjusted p-value of >0.1. **(B)** Volcano plot analysis of differentially expressed genes in day 11 samples. Red dots are upregulated genes, and blue dots are downregulated. Labelled genes are genes in common with the Limb Development term (GO: 0060173). Differentially expressed genes were analysed at a threshold of >1log2 and <-1log2 fold change and an adjusted p-value of >0.1. **(C)** Heat-map visualising representative gene for lineages key gene markers. Red means upregulation, whereas blue means downregulation. Data presented as relative expression to the mean. Only genes with an adjusted p-value<0.1 were considered. **(D)** Visualisation of the significant genes in day 11 samples matching the Cartilage Development term (GO:0051216). Red bars represent upregulated genes, whereas blue bars represent downregulated genes.

### Cells differentiated with the RAPID-E protocol can generate cartilage-like matrix *in vitro*

To evaluate the capacity of cells differentiated with the RAPID-E protocol to generate cartilage-like matrix, we subjected day 11 cells to a further 28 days in pellet culture. Initially, we treated cells with TTNPB for a full 11 days before pellet culture. However, we found that cells would either not spontaneously form pellets at all or the aggregates disintegrated after ∼7 days. Therefore, we removed TTNPB from the differentiation medium at Day 10, thus cells were not treated with TTNPB for 1 day prior to pelleting. Using this approach, cells would form pellets approximately 3 days after centrifugation and these were still compact and spherical at 28 days. Gene expression analyses indicated that cells continued to express chondrogenic genes, with significant upregulation of *SOX9*, *SOX5*, *COL2A1*, *ACAN* and *BMPR1B* (**Figure 5A**). However, pellets also exhibited expression of the fibrotic/hypertrophic-related gene *RUNX2*. Additionally, transcript for *COL10A1*, a marker of pre-hypertrophy, and the hyaline cartilage protein PRG4, were measurable but very low at D28. Histological analyses of pellets at day 28 of culture indicated the presence of sulfated glycosaminoglycans, with pellets stained for Alcian blue, along with a dense collagen matrix, visualised through Picosirius red staining and polarised light microscopy (**Figure 5B**). Immunohistological analyses indicated that cells generated a cartilaginous matrix containing type II collagen and aggrecan, which was sometimes patchy, with negligible expression of type I collagen. Cells within the centre of the pellet expressed SOX9 and the matrix surrounding these cells stained strongest for type II collagen. However, pellets also stained for type X collagen, indicated that cells were beginning to undergo hypertrophy.

**Figure 5:**
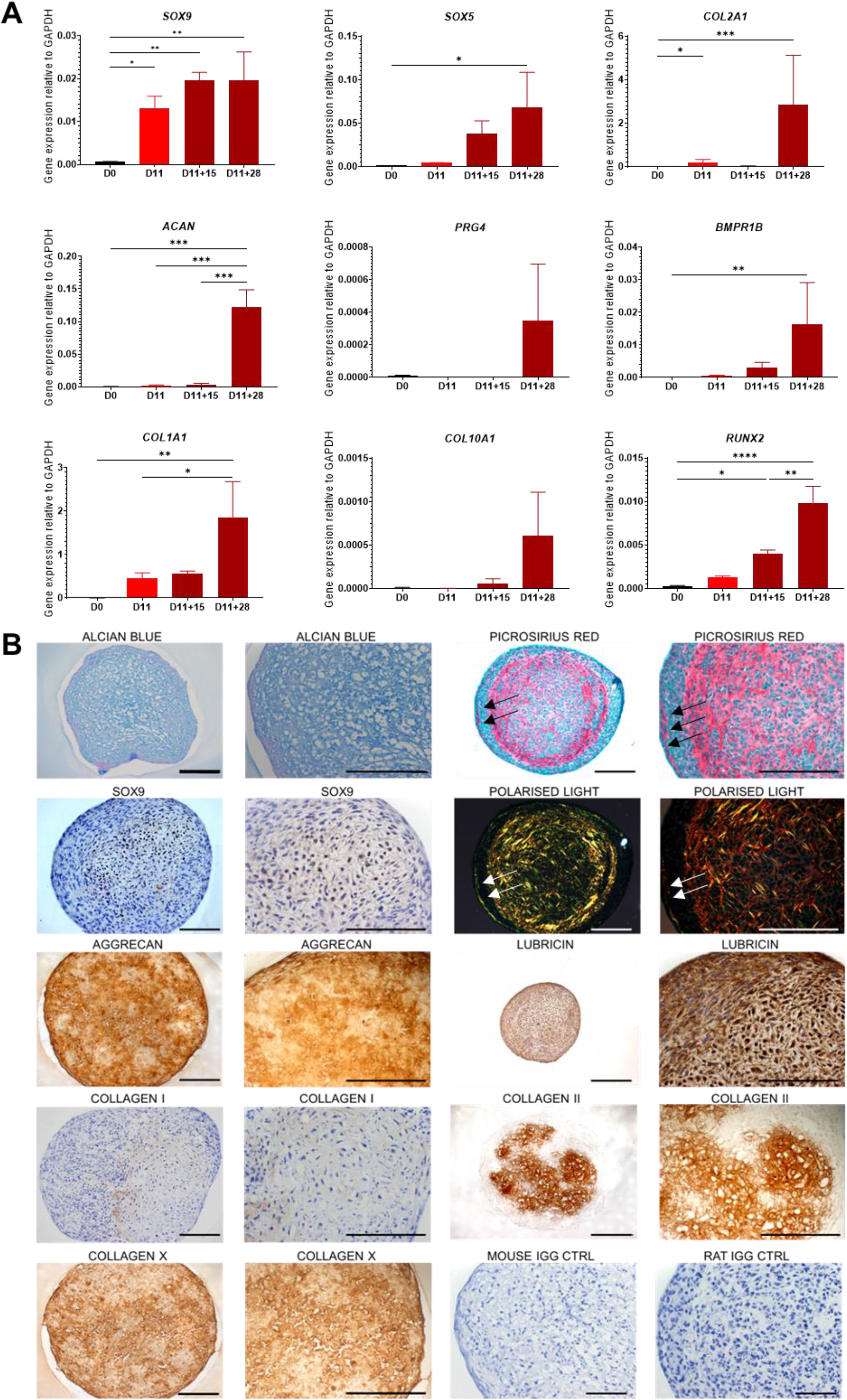
Gene expression analysis and immunohistochemical analysis for chondrogenic markers of RAPID-E cartilaginous pellets. **(A)** RT-qPCR analysis of chondrogenic markers expression at different stages of differentiation in 2D and 3D. Gene expression levels were normalised to the housekeeping gene *GAPDH*. Data are presented as the mean value ± SEM. N numbers for independent experiments in 2D samples (D0 and D11): N=6 for *SOX9, COL2A1, COL1A1, RUNX2*; N=5 *COL10A1*; N=4 for *ACAN* and *BMPR1B*; N=3 *SOX5*; N=2 *PRG4*. N numbers for independent experiments in 3D samples (D11+15 and D11+28): N=4. Statistical analysis was performed using an Ordinary one-way ANOVA test. (* p<0.05, ** p<0.01, *** p<0.005, **** p < 0.0001). **(B)** Histological analysis of pellets protein expression and GAG production. Nuclei were counterstained with haematoxylin (when performed). Secondary antibody control Mouse IgG and Rat IgG are shown. Images representative of N=3 independent experiments. The second and fourth columns show the respective images to the left at higher magnification. Scale bars: 200 µm.

## Discussion

Chondrogenic differentiation of hPSCs has been proposed as a regenerative medicine approach for the treatment of degenerated cartilage in OA. However, the efficiency and reproducibility of hPSC differentiation towards chondrocytes is still limited. We aimed to enhance the developmental relevance and chondrogenic output of our limb bud directed chondrogenic differentiation protocol through activating RAR signalling and modulating the timing of BMP supplementation. We initially attempted to incorporate TTNPB addition into our existing RAPID protocol in the absence of BMP2 addition. BMP signalling is essential for mesodermal and chondrogenic specification during development, enhancing mesenchymal proliferation, condensation and production of cartilage ECM (37,38). Previous studies have indicated that early supplementation of BMP growth factors is required for induction of hPSCs to lateral-plate mesodermal-like a state (6,7,39), whilst inhibition of BMP signals leads to paraxial mesoderm formation (7,40–42). In recent years, other groups have demonstrated that stimulation of hPSCs with only CHIR99021 and TTNPB can efficiently induce mesodermal lineages, as the combination of both molecules induces expression of native BMP ligands, particularly BMP4 which is active during mesodermal specification (19,43–45). Similarly, we observed significant upregulation of *BMP1*, *BMP4* and *BMP5* by day 6 of differentiation, which likely enabled the specification of lateral plate mesodermal lineages. However, the lack of controlled BMP pathway activation through direct stimulation can lead to population heterogeneity of both paraxial and lateral-plate mesodermal cells, as acknowledged by Kawata and colleagues (19).

RNAseq analyses of RAPID-E differentiated cells at day 6, at which time they should exhibit a limb bud mesenchymal-like phenotype, indicated a lack of expression of notable paraxial mesoderm markers, including *MSGN1* and *TBX6* (46,47), whilst lateral plate and limb bud markers, including *FOXF1*, *HAND1* and *PRRX1*, were significantly upregulated. According to our previous findings (6) and the work of Loh and colleagues (6,7), we continued supplementation of additional factors that contribute to restriction of unwanted lineages and specification of mesoderm, in addition to TTNPB addition. For example, RA signalling (48) and also Activin A, which we have used in early mesendoderm induction (16,49), induce reduction of H3K27me3 levels in hESCs (50), which likely establish an appropriate chromatin state for the activation of mesendodermal gene transcription (16,49). Subsequently, we found at day 6 of differentiation that over 70% of the cell population were PRRX1 positive, a key marker of limb bud mesenchyme (51). Thus, with continued stepwise adjustment of growth factor and small molecule combinations we were likely able to maintain induction of a primarily limb bud mesenchymal population even in the absence of BMP growth factor supplementation.

RA signalling plays an important role in skeletal tissue development, first regionalising the lateral plate mesoderm in the posterior and then guiding the forelimb formation process (52–55). Post-cellular ingress, signal transduction involves binding of RA to a RAR, which then forms a heterodimeric complex with retinoid X receptor (RXR). RAR-RXR complexes then directly modulate transcription by binding to RA response elements (RAREs) within the enhancer regions of target genes (54). In the absence of RA, RAR-RXR complexes act as transcriptional repressors, recruiting histone deacetylase (HDAC) and polycomb repressive complex 2 (PRC2) complexes, resulting in chromatin condensation and silencing (56). Cells treated with TTNPB exhibited greater enrichment of biological process terms associated with limb and skeletal development, with a variety of *HOX* genes significantly upregulated. The kinetics of *HOX* gene expression during development is correlated to their response to retinoic acid (57–59), with *HoxA* and *HoxB* clusters displaying a more rapid and stronger response to retinoic acid than *HoxC* and *HoxD* clusters (60). Accordingly, RAPID-E cells exhibited high expression of limb-related *HOX* genes *HOXA9* and *HOXB9* (61) at day 6, whilst at day 11 more posterior *HOX* genes, *HOXC4-6*, *HOXC9* and *HOXD9*, which contribute to distal outgrowth after the formation of the apical ectodermal ridge (62), exhibited their highest expression. Supplementation of retinoic acid has been reported to increase the expression of forelimb marker *TBX5* in mouse ESC-derived limb bud cells (61). However, compared to non-treated cells, TTNPB treated cells at day 6 exhibited lower expression of *TBX5* and higher expression for hind limb marker *TBX4*. By day 11, TTNPB-treated cells exhibited high expression of *TBX5* (similar to mouse ESC data) together with other HOX genes related to forelimb and did not express *TBX4*. This, along with the observed kinetics of *HOX* gene upregulation, suggests that retinoic acid polarisation of differentiation into forelimb takes longer than 2 days to occur. This suggests that a longer period for the limb induction in the RAPID-E protocol could improve the differentiation of cells into limb bud and consequently into chondroprogenitors, as reported by Yamada *et al.* (63).

RA signalling has widely reported roles in a variety of developmental processes, including that of the nervous, pancreatic, neural, cardiac, renal and pulmonary systems (64–68). Some off-target biological GO terms were significantly enriched in samples treated with TTNPB, such as nervous system development (GO:0007399). A recent study demonstrated that addition of 1µM TTNPB can improve the generation of neuronal progenitors from hESCs (69), an effect that should be avoided in the RAPID-E protocol with an addition of only 100nM TTNPB. Furthermore, samples treated at day 11 showed high expression of *RARβ*, which has been showed to negatively regulate neuronal differentiation (69). However, analysis of genes downregulated with addition of TTNPB also revealed an enrichment of terms associated with cardio- and myo-genesis, and expression of cardiac markers *NKX2-5*, *TTNT2, PECAM,* and *TBX20* was decreased, indicating that TTNPB supplementation reduces differentiation towards the heart field, supporting previous literature on retinoic acid signalling (52,70). The restriction of the heart field by TTNPB supplementation may contribute to the channelling of differentiation towards cell populations more relevant to cartilage development.

Activation of retinoic acid signalling to drive chondrogenesis is controversial: its addition has been reported to suppress *SOX9* and cartilage matrix synthesis in chondrocyte cultures (21,71–74). Indeed, we found that removal of TTNPB from growth media was required to enable pellet formation at the end of the adherent phase of the RAPID-E protocol. However, other studies have reported that retinoic acid plays essential roles in limb bud formation and chondrogenesis during skeletal development (53,55,75,76). Kawata and colleagues successfully induced hPSC-chondrogenic differentiated using only CHIR99021 and TTNPB, demonstrating through ATAC-seq and CHIP-seq the important roles of RARα in binding to enhancers of chondrogenic genes *SOX9, SOX5, SOX6* (19). Similarly, we found that addition of TTNPB significantly enhanced *SOX9* expression alongside other associated chondrogenic genes, such as *COL2A1* and *COL11A2* at day 11. TTNPB-treated cells in adherent culture did not express known markers of hypertrophy and the growth plate, including *COL10A1*, *IHH* or *MMP13* and exhibited significantly lower expression of *RUNX2, PTH1R*, *FGFR3* and *VEGFA* in comparison to non-treated samples (77,78). This suggests that, at this stage, retinoic acid signalling had an effect in limiting hypertrophic cartilage differentiation.

Although we found that addition of TTNPB alone significantly enhanced expression of chondrogenic genes by day 6 of differentiation, maintenance of *COL2A1* expression until day 11 was dependent upon BMP2 addition (in combination with GDF5). BMP signalling is known to drive expression of *COL2A1* (36,79) alongside other key chondrogenic genes (17,35,80– 84). Our findings are consistent with those of Yamada and colleagues who reported that stimulation of limb bud mesenchymal-like cells with BMP4 (alongside GDF5 and TGFβ1) was required to generate chondrogenic nodules (63). That BMP2 stimulation at day 9 and 10 enhanced chondrogenesis may be attributed to the high expression of *SOX9* and low expression of *RUNX2*, which together limit BMP2-induced osteogenic differentiation while promoting chondrogenesis (85–88). All the same, further refinement may be needed to avoid the evidence of hypertrophy in the chondrogenic pellets generated from RAPID-E cells. Hypertrophy likely occurred as a result of continued addition of BMP2 throughout the 3D phase, which we showed to have time dependent effects on expression of hypertrophic genes. Addition of ascorbic acid has been reported to increase *COL10A1* expression in chondrogenic ATDC5 cells (89) and chick chondrocytes (90). A recent study, ceased ascorbic acid supplementation in micromass pellets destined for AC differentiation while maintaining it for pellets differentiated towards growth plate cartilage (91). In light of this, modification of growth factor composition in later stages will be required to maintain a non-hypertrophic phenotype. It is worth noting that by immunohistochemistry, pellets showed abundant, although heterogeneous, type II collagen, lubricin and aggrecan expression, suggesting the presence of an AC-committed cell population.

In summary, the RAPID-E protocol represents a promising approach for the differentiation of hPSCs into chondrogenic cells for regenerative medicine applications. Its capability for differentiating cells into chondroprogenitors and its robustness across different hPSC lines make it a valuable tool for future research and potential therapeutic use. Further enrichment approaches may be required to refine the cell population generated in this protocol, allowing for the selection of a more homogeneous articular cartilage progenitor population and cartilage tissue.

## Acknowledgements

Acknowledgements and funding

We thank the following for funding: UKRI EPSRC/MRC Centre for Doctoral training in Regenerative Medicine: Fabrizio E. Mancini, Susan J. Kimber EP/L014904/1; BBSRC: Paul E.A. Humphreys BB/W013940/1, Arthritis Research UK: Susan J. Kimber R20786; MRC UKRMP: Susan J. Kimber MR/K026666; UKRI MRC: Susan J. Kimber, Steven Woods MR/X002020/1; UKRI MRC: Susan J. Kimber MR/S002553/1. Great Ormond Street Hospital (GOSH): Sara Cuvertino. MRC UKRI Doctoral Training Programme: Julieta O’ Flaherty MR/N013751/11. Marco Domingos EP/R00 6 61X/1, EP/S019367/1, EP/P025021/1 and EP/P025498/1. We would like to thank Dr Leo Zeef for the bioinformatic analysis of the RNAseq data and Marie O’Brian for the processing and cutting of the histological samples and the HipSci consortium for iPSC cell line (IERP4, OAPD4 and WIGW2).

## Author Contributions

F.E.M., S.J.K, P.E.A.H and S.W designed the experiments and provided input for experimental analyses and data interpretation. F.E.M. and P.E.A.H wrote the main manuscript and prepared the figures. F.E.M. conducted all the main experiments reported here. N.B performed histological staining of 3D pellet samples. S.C performed the differentiation by RAPID-E on hiPSCs (WIGW2, JF191B, IERP4 and OAPD4). J.OF. generated the JF191B hiPSC line. S.W generated the SW163D hiPSC line. S.J.K., M.A.N.D. and L.B supervised the project. All authors reviewed and approved the final manuscript.

## Competing interests

The authors declare no competing interests.

## Notes

### Competing Interest Statement

The authors have declared no competing interest.

## References

1. James, S. L., Abate, D., Abate, K. H., Abay, S. M., Abbafati, C., Abbasi, N., Abbastabar, H., Abd-Allah, F., Abdela, J., Abdelalim, A., Abdollahpour, I., Abdulkader, R. S., Abebe, Z., Abera, S. F., Abil, O. Z., Abraha, H. N., Abu-Raddad, L. J., Abu-Rmeileh, N. M. E., Accrombessi, M. M. K. et al. Global, regional, and national incidence, prevalence, and years lived with disability for 354 Diseases and Injuries for 195 countries and territories, 1990-2017: A systematic analysis for the Global Burden of Disease Study 2017. The Lancet 392, 1789–1858 (2018).

2. Zhang, W., Ouyang, H., Dass, C. R. & Xu, J. Current research on pharmacologic and regenerative therapies for osteoarthritis. Bone Research 4, 15040 (2016).

3. Yamanaka, S. Pluripotent Stem Cell-Based Cell Therapy—Promise and Challenges. Cell Stem Cell 27, 523–531 (2020).

4. Jevotovsky, D. S., Alfonso, A. R., Einhorn, T. A. & Chiu, E. S. Osteoarthritis and stem cell therapy in humans: a systematic review. Osteoarthritis and Cartilage 26, 711–729 (2018).

5. Craft, A. M., Rockel, J. S., Nartiss, Y., Kandel, R. A., Alman, B. A. & Keller, G. M. Generation of articular chondrocytes from human pluripotent stem cells. Nature Biotechnology 33, 638–645 (2015).

6. Smith, C. A., Humphreys, P. A., Naven, M. A., Woods, S., Mancini, F. E., O’Flaherty, J., Meng, Q.-J. & Kimber, S. J. Directed differentiation of hPSCs through a simplified lateral plate mesoderm protocol for generation of articular cartilage progenitors. Plos One 18, e0280024 (2023).

7. Loh, K. M. M., Chen, A., Koh, P. W. W., Deng, T. Z. Z., Sinha, R., Tsai, J. M. M., Barkal, A. A. A., Shen, K. Y. Y., Jain, R., Morganti, R. M. M., Shyh-Chang, N., Fernhoff, N. B. B., George, B. M. M., Wernig, G., Salomon, R. E. E. A., Chen, Z., Vogel, H., Epstein, J. A. A., Kundaje, A. et al. Mapping the Pairwise Choices Leading from Pluripotency to Human Bone, Heart, and Other Mesoderm Cell Types. Cell 166, 451–467 (2016).

8. Chijimatsu, R. & Saito, T. Mechanisms of synovial joint and articular cartilage development. Cellular and Molecular Life Sciences 76, 3939–3952 (2019).

9. Humphreys, P. A., Mancini, F. E., Ferreira, M. J. S., Woods, S., Ogene, L. & Kimber, S. J. Developmental principles informing human pluripotent stem cell differentiation to cartilage and bone. Seminars in Cell and Developmental Biology 127, 17–36 (2022).

10. Sumi, T., Tsuneyoshi, N., Nakatsuji, N. & Suemori, H. Defining early lineage specification of human embryonic stem cells by the orchestrated balance canonical Wnt/β-catenin, activin/Nodal and BMP signaling. Development 135, 2969–2979 (2008).

11. Row, R. H., Pegg, A., Kinney, B. A., Farr, G. H., Maves, L., Lowell, S., Wilson, V. & Martin, B. L. BMP and FGF signaling interact to pattern mesoderm by controlling basic helix-loop-helix transcription factor activity. eLife 7, 1–27 (2018).

12. Pignatti, E., Zeller, R. & Zuniga, A. To BMP or not to BMP during vertebrate limb bud development. Seminars in Cell and Developmental Biology 32, 119–127 (2014).

13. Ray, A., Singh, P. N. P., Sohaskey, M. L., Harland, R. M. & Bandyopadhyay, A. Precise spatial restriction of BMP signaling is essential for articular cartilage differentiation. Development 142, 1169–1179 (2015).

14. Kobayashi, T., Lyons, K. M., McMahon, A. P. & Kronenberg, H. M. BMP signaling stimulates cellular differentiation at multiple steps during cartilage development. Proceedings of the National Academy of Sciences of the United States of America 102, 18023–18027 (2005).

15. Yoon, B. S. & Lyons, K. M. Multiple functions of BMPs in chondrogenesis. Journal of Cellular Biochemistry 93, 93–103 (2004).

16. Oldershaw, R. A., Baxter, M. A., Lowe, E. T., Bates, N., Grady, L. M., Soncin, F., Brison, D. R., Hardingham, T. E. & Kimber, S. J. Directed differentiation of human embryonic stem cells toward chondrocytes. Nature Biotechnology 28, 1187–1194 (2010).

17. Wang, T., Nimkingratana, P., Smith, C. A., Cheng, A., Hardingham, T. E. & Kimber, S. J. Enhanced chondrogenesis from human embryonic stem cells. Stem Cell Research 39, 101497 (2019).

18. Diederichs, S., Klampfleuthner, F. A. M., Moradi, B. & Richter, W. Chondral Differentiation of Induced Pluripotent Stem Cells Without Progression Into the Endochondral Pathway. Frontiers in Cell and Developmental Biology 7, 1–10 (2019).

19. Kawata, M., Mori, D., Kanke, K., Hojo, H., Ohba, S., Chung, U. il, Yano, F., Masaki, H., Otsu, M., Nakauchi, H., Tanaka, S. & Saito, T. Simple and Robust Differentiation of Human Pluripotent Stem Cells toward Chondrocytes by Two Small-Molecule Compounds. Stem Cell Reports 13, 530–544 (2019).

20. Weston, A. D., Chandraratna, R. A. S., Torchia, J. & Underhill, T. M. Requirement for RAR-mediated gene repression in skeletal progenitor differentiation. Journal of Cell Biology 158, 39–51 (2002).

21. Pacifici, M., Cossu, G., Molinaro, M. & Tato, F. Vitamin A inhibits chondrogenesis but not myogenesis. Experimental Cell Research 129, 469–474 (1980).

22. Hoffman, L. M., Garcha, K., Karamboulas, K., Cowan, M. F., Drysdale, L. M., Horton, W. A. & Underhill, T. M. BMP action in skeletogenesis involves attenuation of retinoid signaling. Journal of Cell Biology 174, 101–113 (2006).

23. Langston, A. W. & Gudas, L. J. Retinoic acid and homeobox gene regulation. Current Opinion in Genetics and Development 4, 550–555 (1994).

24. Boncinelli, E., Simeone, A., Acampora, D. & Mavilio, F. HOX gene activation by retinoic acid. Trends in Genetics 7, 329–334 (1991).

25. Bel-Vialar, S., Itasaki, N. & Krumlauf, R. Initiating Hox gene expression: In the early chick neural tube differential sensitivity to FGF and RA signaling subdivides the HoxB genes in two distinct groups. Development 129, 5103–5115 (2002).

26. Ye, J., Coulouris, G., Zaretskaya, I., Cutcutache, I., Rozen, S. & Madden, T. L. Primer-BLAST: a tool to design target-specific primers for polymerase chain reaction. BMC bioinformatics 13, 134 (2012).

27. Xie, Z., Bailey, A., Kuleshov, M. V., Clarke, D. J. B., Evangelista, J. E., Jenkins, S. L., Lachmann, A., Wojciechowicz, M. L., Kropiwnicki, E., Jagodnik, K. M., Jeon, M. & Ma’ayan, A. Gene Set Knowledge Discovery with EnrichR. Current Protocols 1, e90 (2021).

28. Kuleshov, M. V, Jones, M. R., Rouillard, A. D., Fernandez, N. F., Duan, Q., Wang, Z., Koplev, S., Jenkins, S. L., Jagodnik, K. M., Lachmann, A., Mcdermott, M. G., Monteiro, C. D., Gundersen, W. & Ma, A. EnrichR: a comprehensive gene set enrichment analysis web server 2016 update. Nucleic Acids Research 44, 90–97 (2016).

29. Chen, E., Tan, C., Kou, Y., Duan, Q., Wang, Q., Meirelles, G., Clark, N. & Ma’ayan, A. EnrichR: interactive and collaborative HTML5 gene list enrichment analysis tool. BMC Bioinformatics 14, 1–14 (2013).

30. Goedhart, J. & Luijsterburg, M. S. VolcaNoseR is a web app for creating, exploring, labeling and sharing volcano plots. Scientific Reports 2020 10:1 10, 1–5 (2020).

31. Eisen, M. B., Spellman, P. T., Brown, P. O. & Botstein, D. Cluster analysis and display of genome-wide expression patterns. Proceedings of the National Academy of Sciences of the United States of America 95, 14863–14868 (1998).

32. Humphreys, P. A., Woods, S., Smith, C. A., Bates, N., Cain, S. A., Lucas, R. & Kimber, S. J. Optogenetic control of the BMP signaling pathway. ACS Synthetic Biology 9, 3067–3078 (2020).

33. Shen, P., Chen, L., Zhang, D., Xia, S., Lv, Z., Zou, D., Zhang, Z., Yang, C. & Li, W. Rapid induction and long-term self-renewal of neural crest-derived ectodermal chondrogenic cells from hPSCs. npj Regenerative Medicine 7, 1–15 (2022).

34. Xu, S. C., Harris, M. A., Rubenstein, J. L. R., Mundy, G. R. & Harris, S. E. Bone morphogenetic protein-2 (BMP-2) signaling to the Col2α1 gene in chondroblasts requires the homeobox gene Dlx-2. DNA and Cell Biology 20, 359–365 (2001).

35. Lafont, J. E., Poujade, F. A., Pasdeloup, M., Neyret, P. & Mallein-Gerin, F. Hypoxia potentiates the BMP-2 driven COL2A1 stimulation in human articular chondrocytes via p38 MAPK. Osteoarthritis and Cartilage 24, 856–867 (2016).

36. Fernández-Lloris, R., Viñals, F., López-Rovira, T., Harley, V., Bartrons, R., Rosa, J. L. & Ventura, F. Induction of the Sry-Related Factor SOX6 Contributes to Bone Morphogenetic Protein-2-Induced Chondroblastic Differentiation of C3H10T1/2 Cells. Molecular Endocrinology 17, 1332–1343 (2003).

37. Kim, H., Neugebauer, J., Mcknite, A., Tilak, A. & Christian, J. L. BMP7 functions predominantly as a heterodimer with BMP2 or BMP4 during mammalian embryogenesis. 1–22 (2019).

38. James, R. G. & Schultheiss, T. M. Bmp signaling promotes intermediate mesoderm gene expression in a dose-dependent, cell-autonomous and translation-dependent manner. Developmental Biology 288, 113–125 (2005).

39. Huang, D., Li, J., Hu, F., Xia, C., Weng, Q., Wang, T., Peng, H., Wu, B., Wu, H., Xiong, J., Lin, Y., Wang, Y., Zhang, Q., Liu, X., Liu, L., Zheng, X., Geng, Y., Du, X., Zhu, X. et al. Lateral plate mesoderm cell-based organoid system for NK cell regeneration from human pluripotent stem cells. Cell Discovery 8, (2022).

40. Xi, H., Fujiwara, W., Gonzalez, K., Jan, M., Liebscher, S., Van Handel, B., Schenke-Layland, K. & Pyle, A. D. In Vivo Human Somitogenesis Guides Somite Development from hPSCs. Cell Reports 18, 1573–1585 (2017).

41. Wu, C., Dicks, A., Steward, N., Tang, R., Katz, D. B., Choi, Y. & Guilak, F. Single cell transcriptomic analysis of human pluripotent stem cell chondrogenesis. Nature Communications 12, 362 (2021).

42. Umeda, K., Zhao, J., Simmons, P., Stanley, E., Elefanty, A. & Nakayama, N. Human chondrogenic paraxial mesoderm, directed specification and prospective isolation from pluripotent stem cells. Scientific Reports 2, 1–11 (2012).

43. Araoka, T., Mae, S. I., Kurose, Y., Uesugi, M., Ohta, A., Yamanaka, S. & Osafune, K. Efficient and rapid induction of human iPSCs/ESCs into nephrogenic intermediate mesoderm using small molecule-based differentiation methods. PLoS ONE 9, epub (1-14) (2014).

44. Zhang, P., Li, J., Tan, Z., Wang, C., Liu, T., Chen, L., Yong, J., Jiang, W., Sun, X., Du, L., Ding, M. & Deng, H. Short-term BMP-4 treatment initiates mesoderm induction in human embryonic stem cells. Blood 111, 1933–1941 (2008).

45. Winnier, G., Blessing, M., Labosky, P. A. & Hogan, B. L. M. Bone morphogenetic protein-4 is required for mesoderm formation and patterning in the mouse. Genes and Development 9, 2105–2116 (1995).

46. Chalamalasetty, R. B., Garriock, R. J., Dunty, W. C., Kennedy, M. W., Jailwala, P., Si, H. & Yamaguchi, T. P. Mesogenin 1 is a master regulator of paraxial presomitic mesoderm differentiation. Development (Cambridge) 141, 4285–4297 (2014).

47. Chapman, D. L., Agulnik, I., Hancock, S., Silver, L. M. & Papaioannou, V. E. Tbx6, a mouse T-box gene implicated in paraxial mesoderm formation at gastrulation. Developmental Biology 180, 534–542 (1996).

48. Kashyap, V. & Gudas, L. J. Epigenetic regulatory mechanisms distinguish retinoic acid-mediated transcriptional responses in stem cells and fibroblasts. Journal of Biological Chemistry 285, 14534–14548 (2010).

49. Cheng, A., Kapacee, Z., Peng, J., Lu, S., Lucas, R. J., Hardingham, T. E. & Kimber, S. J. Cartilage Repair Using Human Embryonic Stem Cell-Derived Chondroprogenitors. Stem Cells Translational Medicine 3, 1287–1294 (2014).

50. Wang, L., Xu, X., Cao, Y., Li, Z., Cheng, H., Zhu, G., Duan, F., Na, J., Han, J. D. J. & Chen, Y. G. Activin/Smad2-induced histone H3 Lys-27 trimethylation (H3K27me3) reduction is crucial to initiate mesendoderm differentiation of human embryonic stem cells. Journal of Biological Chemistry 292, 1339–1350 (2017).

51. Fowler, D. A. & Larsson, H. C. E. The tissues and regulatory pattern of limb chondrogenesis. Developmental Biology 463, 124–134 (2020).

52. Waxman, J. S., Keegan, B. R., Roberts, R. W., Poss, K. D. & Yelon, D. Hoxb5b Acts Downstream of Retinoic Acid Signaling in the Forelimb Field to Restrict Heart Field Potential in Zebrafish. Developmental Cell 15, 923–934 (2008).

53. Feneck, E. & Logan, M. The role of retinoic acid in establishing the early limb bud. Biomolecules 10, (2020).

54. Cunningham, T. J. & Duester, G. Mechanisms of retinoic acid signalling and its roles in organ and limb development. Nature Reviews Molecular Cell Biology 16, 110–123 (2015).

55. Nishimoto, S., Wilde, S. M., Wood, S. & Logan, M. P. O. RA Acts in a Coherent Feed-Forward Mechanism with Tbx5 to Control Limb Bud Induction and Initiation. Cell Reports 12, 879–891 (2015).

56. Jepsen, K., Hermanson, O., Onami, T. M., Gleiberman, A. S., Lunyak, V., McEvilly, R. J., Kurokawa, R., Kumar, V., Liu, F., Seto, E., Hedrick, S. M., Mandel, G., Glass, C. K., Rose, D. W. & Rosenfeld, M. G. Combinatorial roles of the nuclear receptor corepressor in transcription and development. Cell 102, 753–763 (2000).

57. Simeone, A., Acampora, D., Arcioni, L., Andrews, P. W., Boncinelli, E. & Mavilio, F. Sequential activation of HOX2 homeobox genes by retinoic acid in human embryonal carcinoma cells. Nature 1990 346:6286 346, 763–766 (1990).

58. Kmita, M. & Duboule, D. Organizing Axes in Time and Space; 25 Years of Colinear Tinkering. Science 301, 331–333 (2003).

59. Papalopulu, N., Lovel-badage, R. & Krumlauf, R. The expression of murine Hox-2 genes is dependent on the differentiation pathway and displays a collinear sensitivity to retinoic acid in F9 cells and Xenopus embryos. Nucleic Acids Research 19, 5497 (1991).

60. De Kumar, B., Parrish, M. E., Slaughter, B. D., Unruh, J. R., Gogol, M., Seidel, C., Paulson, A., Li, H., Gaudenz, K., Peak, A., McDowell, W., Fleharty, B., Ahn, Y., Lin, C., Smith, E., Shilatifard, A. & Krumlauf, R. Analysis of dynamic changes in retinoid-induced transcription and epigenetic profiles of murine Hox clusters in ES cells. Genome Research 25, 1229–1243 (2015).

61. Mori, S., Sakakura, E., Tsunekawa, Y., Hagiwara, M., Suzuki, T. & Eiraku, M. Self-organized formation of developing appendages from murine pluripotent stem cells. Nature Communications 10, 1–13 (2019).

62. Sheth, R., Grégoire, D., Dumouchel, A., Scotti, M., Trang Pham, J. M., Nemec, S., Félix Bastida, M., Ros, M. A. & Kmita, M. Decoupling the function of Hox and Shh in developing limb reveals multiple inputs of Hox genes on limb growth. Development 140, 2130–2138 (2013).

63. Yamada, D., Nakamura, M., Takao, T., Takihira, S., Yoshida, A., Kawai, S., Miura, A., Ming, L., Yoshitomi, H., Gozu, M., Okamoto, K., Hojo, H., Kusaka, N., Iwai, R., Nakata, E., Ozaki, T., Toguchida, J. & Takarada, T. Induction and expansion of human PRRX1+ limb-bud-like mesenchymal cells from pluripotent stem cells. Nature Biomedical Engineering 5, 926–940 (2021).

64. Rosselot, C., Spraggon, L., Chia, I., Batourina, E., Riccio, P., Lu, B., Niederreither, K., Dolle, P., Duester, G., Chambon, P., Costantini, F., Gilbert, T., Molotkov, A. & Mendelsohn, C. Non-cell-autonomous retinoid signaling is crucial for renal development. Development 137, 283–292 (2010).

65. Jacobs, S., Lie, D. C., DeCicco, K. L., Shi, Y., DeLuca, L. M., Gage, F. H. & Evans, R. M. Retinoic acid is required early during adult neurogenesis in the dentate gyrus. Proceedings of the National Academy of Sciences of the United States of America 103, 3902–3907 (2006).

66. Wiesinger, A., Boink, G. J. J., Christoffels, V. M. & Devalla, H. D. Retinoic acid signaling in heart development: Application in the differentiation of cardiovascular lineages from human pluripotent stem cells. Stem Cell Reports 16, 2589–2606 (2021).

67. Lorberbaum, D. S., Kishore, S., Rosselot, C., Sarbaugh, D., Brooks, E. P., Aragon, E., Xuan, S., Simon, O., Ghosh, D., Mendelsohn, C., Gadue, P. & Sussel, L. Retinoic acid signaling within pancreatic endocrine progenitors regulates mouse and human β cell specification. Development (Cambridge) 147, (2020).

68. Fernandes-Silva, H., Araújo-Silva, H., Correia-Pinto, J. & Moura, R. S. Retinoic acid: A key regulator of lung development. Biomolecules 10, 1–18 (2020).

69. Das, M. & Pethe, P. Differential expression of retinoic acid alpha and beta receptors in neuronal progenitors generated from human embryonic stem cells in response to TTNPB (a retinoic acid mimetic). Differentiation 121, 13–24 (2021).

70. Duong, T. B., Holowiecki, A. & Waxman, J. S. Retinoic acid signaling restricts the size of the first heart field within the anterior lateral plate mesoderm. Developmental Biology 473, 119–129 (2021).

71. Sumitani, Y., Uchibe, K., Yoshida, K., Weng, Y., Guo, J., Yuan, H., Ikegame, M., Kamioka, H. & Okamura, H. Inhibitory effect of retinoic acid receptor agonists on in vitro chondrogenic differentiation. Anatomical Science International 95, 202–208 (2020).

72. Cho, S. H., Oh, C. Do, Kim, S. J., Kim, I. C. & Chun, J. S. Retinoic acid inhibits chondrogenesis of mesenchymal cells by sustaining expression of N-cadherin and its associated proteins. Journal of Cellular Biochemistry 89, 837–847 (2003).

73. He, N., Brysk, H., Tyring, S. K., Ohkubo, I. & Brysk, M. M. Transcriptional suppression of Sox9 expression in chondrocytes by retinoic acid. Journal of Cellular Biochemistry 81, 71–78 (2001).

74. Pacifici, M. Retinoid roles and action in skeletal development and growth provide the rationale for an ongoing heterotopic ossification prevention trial. Bone 109, 267–275 (2018).

75. Niederreither, K., Subbarayan, V., Dollé, P. & Chambon, P. Embryonic retinoic acid synthesis is essential for early mouse post-implantation development. Nature genetics 21, 444–448 (1999).

76. Niederreither, K., Vermot, J., Schuhbaur, B., Chambon, P. & Dollé, P. Embryonic retinoic acid synthesis is required for forelimb growth and anteroposterior patterning in the mouse. Development 129, 3563–3574 (2002).

77. Zhang, W., Chen, J., Zhang, S. & Ouyang, H. W. Inhibitory function of parathyroid hormone-related protein on chondrocyte hypertrophy: the implication for articular cartilage repair. Arthritis Research and Therapy 14, 1–10 (2012).

78. Oh, C. Do, Lu, Y., Liang, S., Mori-Akiyama, Y., Chen, D., De Crombrugghe, B. & Yasuda, H. SOX9 regulates multiple genes in chondrocytes, including genes encoding ECM proteins, ECM modification enzymes, receptors, and transporters. PLoS ONE 9, 107577 (2014).

79. Timur, U. T., Caron, M., van den Akker, G., van der Windt, A., Visser, J., van Rhijn, L., Weinans, H., Welting, T., Emans, P. & Jahr, H. Increased TGF-β and BMP levels and improved chondrocyte-specific marker expression in vitro under cartilage-specific physiological osmolarity. International Journal of Molecular Sciences 20, (2019).

80. Shu, B., Zhang, M., Xie, R., Wang, M., Jin, H., Hou, W., Tang, D., Harris, S. E., Mishina, Y., O’Keefe, R. J., Hilton, M. J., Wang, Y. & Chen, D. BMP2, but not BMP4, is crucial for chondrocyte proliferation and maturation during endochondral bone development. Journal of Cell Science 124, 3428–3440 (2011).

81. Kramer, J., Hegert, C., Guan, K., Wobus, A. M., Müller, P. K. & Rohwedel, J. Embryonic stem cell-derived chondrogenic differentiation in vitro: activation by BMP-2 and BMP-4. Mechanisms of Development 92, 193–205 (2000).

82. Murphy, M. K., Huey, D. J., Hu, J. C. & Athanasiou, K. A. TGF-β1, GDF-5, and BMP-2 stimulation induces chondrogenesis in expanded human articular chondrocytes and marrow-derived stromal cells. Stem Cells 33, 762–773 (2015).

83. Davidson, E. N. B., Vitters, E. L., van Lent, P. L. E. M., van de Loo, F. A. J., van den Berg, W. B. & van der Kraan, P. M. Elevated extracellular matrix production and degradation upon bone morphogenetic protein-2 (BMP-2) stimulation point toward a role for BMP-2 in cartilage repair and remodeling. Arthritis Research and Therapy 9, 1– 11 (2007).

84. Gründer, T., Gaissmaier, C., Fritz, J., Stoop, R., Hortschansky, P., Mollenhauer, J. & Aicher, W. K. Bone morphogenetic protein (BMP)-2 enhances the expression of type II collagen and aggrecan in chondrocytes embedded in alginate beads. Osteoarthritis and Cartilage 12, 559–567 (2004).

85. Liao, J., Hu, N., Zhou, N., Zhao, C., Liang, X., Chen, H., Xu, W., Chen, C., Cheng, Q. & Huang, W. Sox9 Potentiates BMP2-Induced Chondrogenic Differentiation and Inhibits BMP2-Induced Osteogenic Differentiation. Regenerative Medicine and Plastic Surgery 263–280 (2019). doi:10.1007/978-3-030-19962-3_19

86. Phimphilai, M., Zhao, Z., Boules, H., Roca, H. & Franceschi, R. T. BMP signaling is required for RUNX2-dependent induction of the osteoblast phenotype. Journal of Bone and Mineral Research 21, 637–646 (2006).

87. Zhou, G., Zheng, Q., Engin, F., Munivez, E., Chen, Y., Sebald, E., Krakow, D. & Lee, B. Dominance of SOX9 function over RUNX2 during skeletogenesis. Proceedings of the National Academy of Sciences of the United States of America 103, 19004–19009 (2006).

88. Haseeb, A., Kc, R., Angelozzi, M., de Charleroy, C., Rux, D., Tower, R. J., Yao, L., da Silva, R. P., Pacifici, M., Qin, L. & Lefebvre, V. SOX9 keeps growth plates and articular cartilage healthy by inhibiting chondrocyte dedifferentiation/ osteoblastic redifferentiation. Proceedings of the National Academy of Sciences of the United States of America 118, 1–11 (2021).

89. Kirimoto, A., Takagi, Y., Ohya, K. & Shimokawa, H. Effects of retinoic acid on the differentiation of chondrogenic progenitor cells, ATDC5. Journal of Medical and Dental Sciences 52, 153–162 (2005).

90. Pacifici, M., Golden, E. B., Iwamoto, M. & Adams, S. L. Retinoic acid treatment induces type X collagen gene expression in cultured chick chondrocytes. Experimental Cell Research 195, 38–46 (1991).

91. Richard, D., Pregizer, S., Venkatasubramanian, D., Raftery, R. M., Muthuirulan, P., Liu, Z., Capellini, T. D. & Craft, A. M. Lineage-specific differences and regulatory networks governing human chondrocyte development. 12, 79925 (2023).

92. Fosang, A. J., Tyler, J. A. & Hardingham, T. E. Effect of interleukin-1 and insulin like growth factor-1 on the release of proteoglycan components and hyaluronan from pig articular cartilage in explant culture. Matrix (Stuttgart, Germany) 11, 17–24 (1991).

