## Supplementary Information for "Effect of a retinoic acid analogue on BMP-driven pluripotent stem cell chondrogenesis"

**List of Supporting Information**

### Supplementary Figures

**Supplementary Figure 1:** Quantification of BMP2 batch and manufacturer activity in TC28a2-BRE/P cells.

**Supplementary Figure 2** - Cell proliferation of MAN13 hESCs during differentiation induced by the RAPID-E protocol.

**Supplementary Figure 3** - RT-qPCR analysis of chondrogenic marker expression in hiPSCs differentiated by the RAPID-E protocol.

**Supplementary Figure 4** - Hierarchical clustering analysis of all samples comparing the top ~10,000 most highly expressed genes that exhibited at least 2-fold change between two or more samples.

**Supplementary Figure 5** –Analysis of transcriptional changes induced by TTNPB addition

**Supplementary Figure 6** - Flow cytometry analysis for the limb-bud mesenchyme marker PRRX1 in MAN13 hESCs differentiated by the RAPID-E protocol.

**Supplementary Figure 7** - Analysis of HOX gene expression influenced by TTNPB addition.

**Supplementary Figure 8** - Gene expression of articular and hypertrophic cartilage markers.

### Supplementary Tables

**Supplementary Table 1 –** Overview of the RAPID-E differentiation protocol – For allowing pellet formation, TTNPB supplementation was ceased at day 10.

**Supplementary Table 2** - Overview of antibodies used for immunohistochemistry and the concentration used.

**Supplementary Table 3 –** Primers used for RT-qPCR

### Supplementary Figures

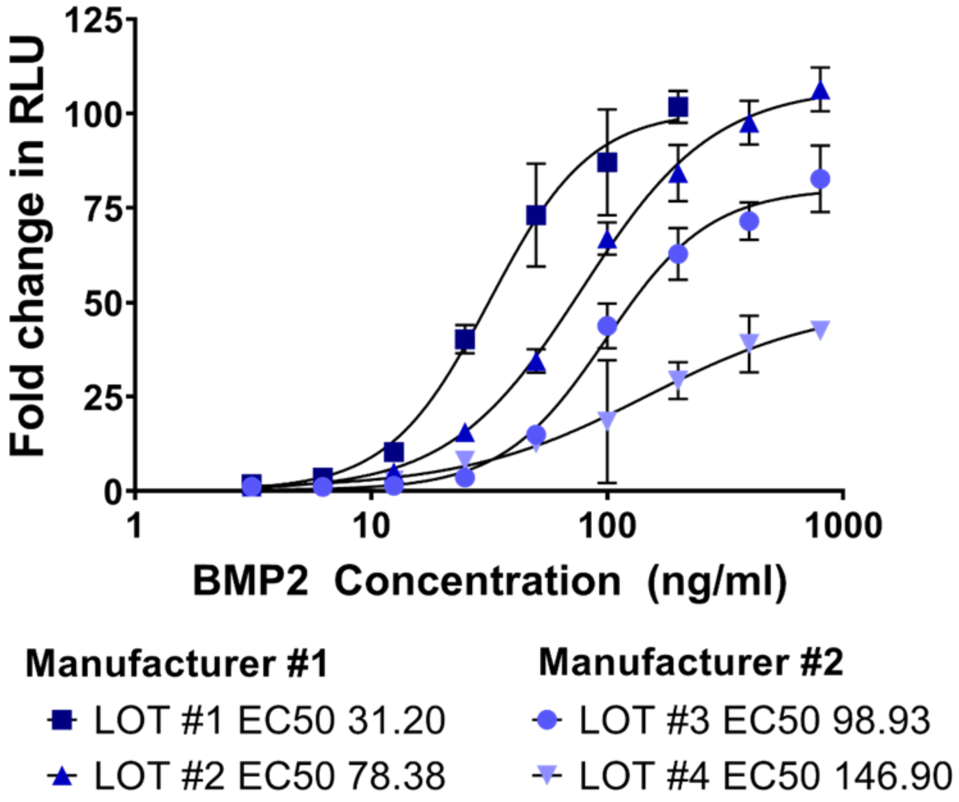


**Supplementary Figure 1 - Quantification of BMP2 batch and manufacturer activity in TC28a2-BRE/P cells.** Data presented as mean fold change in RLU from unstimulated cells ± SD. n = 3 wells per condition.

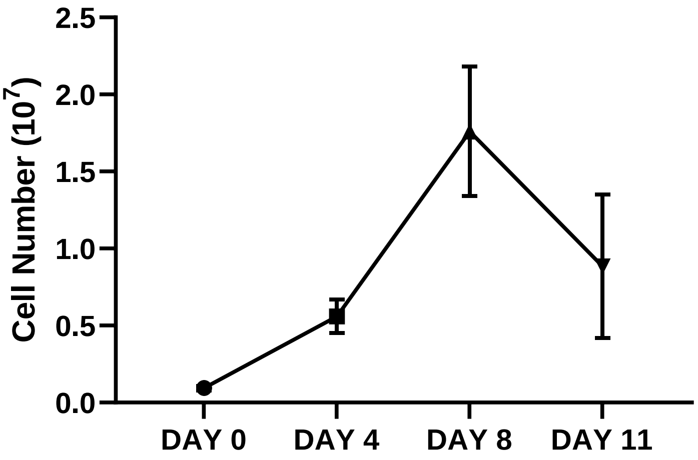


**Supplementary Figure 2 - Cell proliferation of MAN13 hESCs during differentiation induced by the RAPID-E protocol.** Data are presented as means ± SEM from N=2 independent experiments.

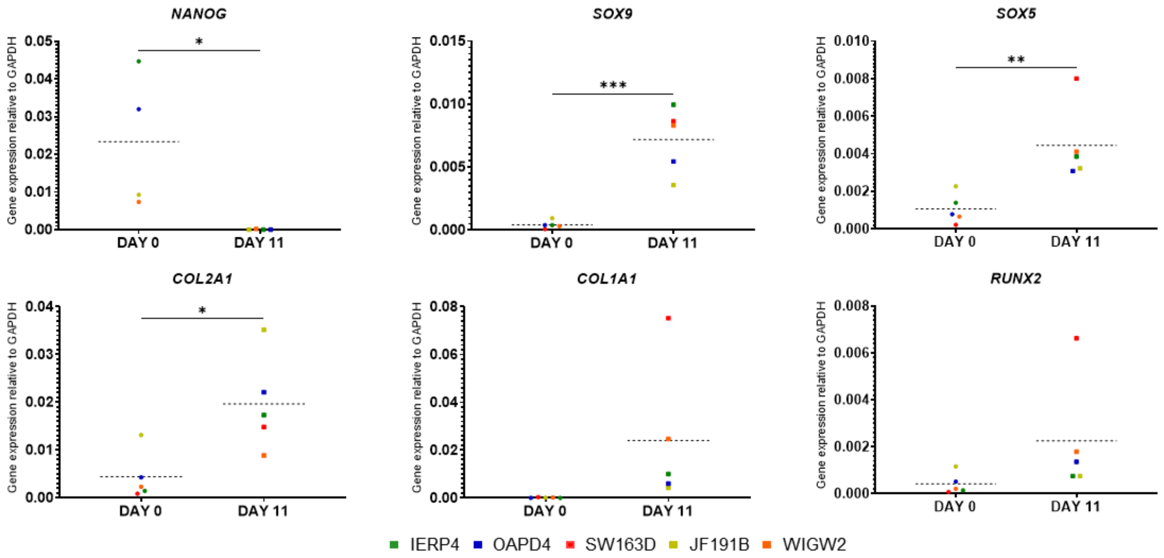


**Supplementary Figure 3 - RT-qPCR analysis of chondrogenic marker expression in hiPSCs differentiated by the RAPID-E protocol.** Data presented as individual values of 5 independent biological repeats using different hiPSCs lines: IERP4, OAPD4, SW163D, JF191B, and WIGW2. The dotted horizontal line represents the mean value. Statistical significance was calculated using an unpaired t-test (p<0.05 *. p<0.01 **, p<0.0005 ***).

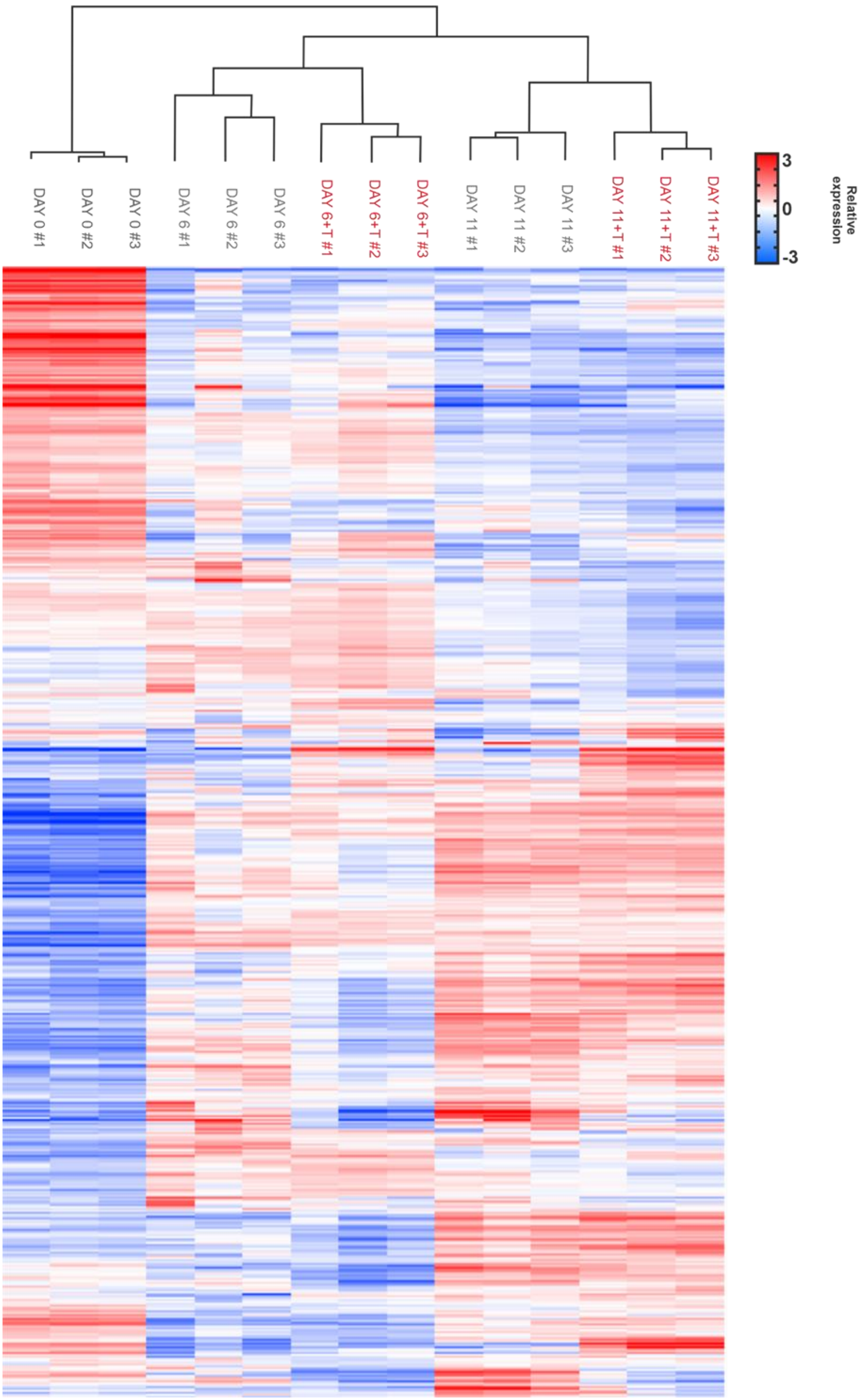


**Supplementary Figure 4 -** Hierarchical clustering analysis of all samples comparing the top ~10,000 most highly expressed genes that exhibited at least 2-fold change between two or more samples. Clustering and visualisation performed using Cluster 3.0 and Treeview software (31).

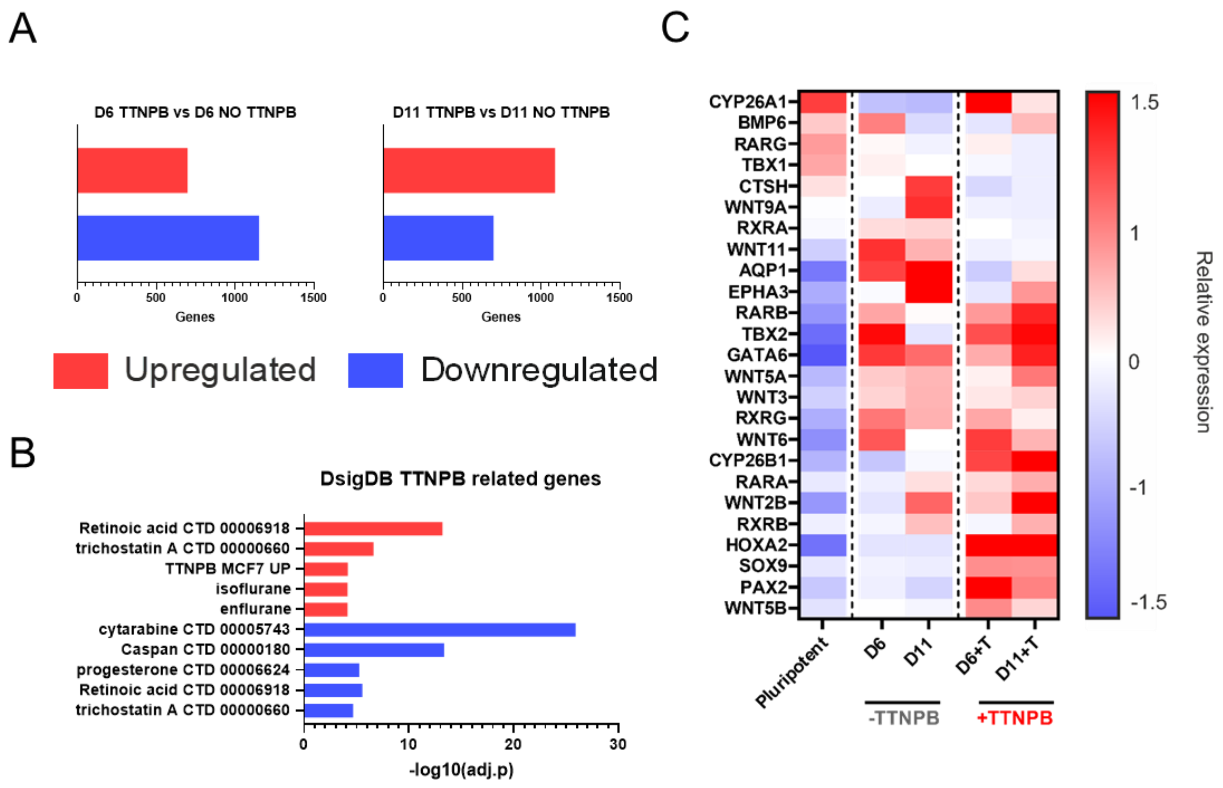


**Supplementary Figure 5 –Analysis of transcriptional changes induced by TTNPB addition. (A)** Histograms indicating number of differentially expressed genes when comparing Day 6 or Day 11 samples differentiated with or without TTNPB addition. **(B)** Dsig enrichment analysis of common genes between Day 11 and day 6 samples. **(C)** Heatmap illustrating expression profiles of genes regulated by retinoic acid signalling**.** The data are presented as relative expression to the mean, with upregulation represented by red and downregulation by blue. Only genes with an adjusted p-value <0.1 were included in the heatmap.

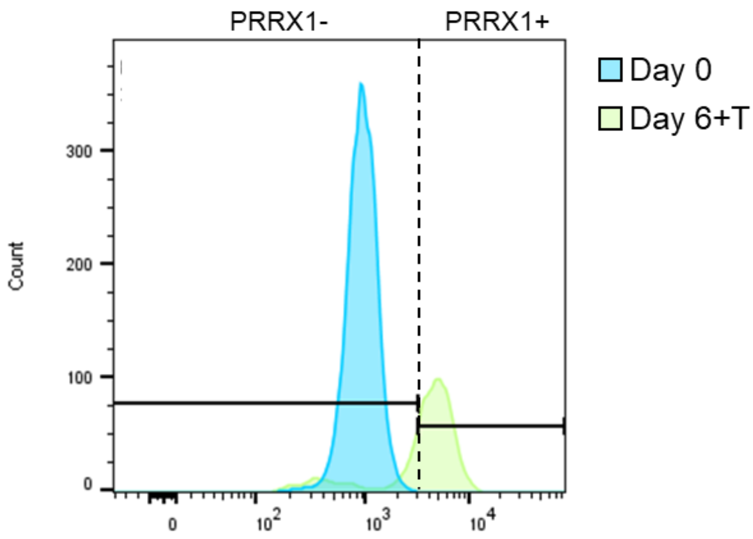


**Supplementary Figure 6 - Flow cytometry analysis for the limb-bud mesenchyme marker PRRX1 in MAN13 hESCs differentiated by the RAPID-E protocol.** The dotted line represents the gate threshold, dividing the population into positive and negative for the marker considered. Light Blue histogram represent M13 hESCs, the green histogram represent M13 hESCs differentiated by the RAPID-E protocol at day 6.

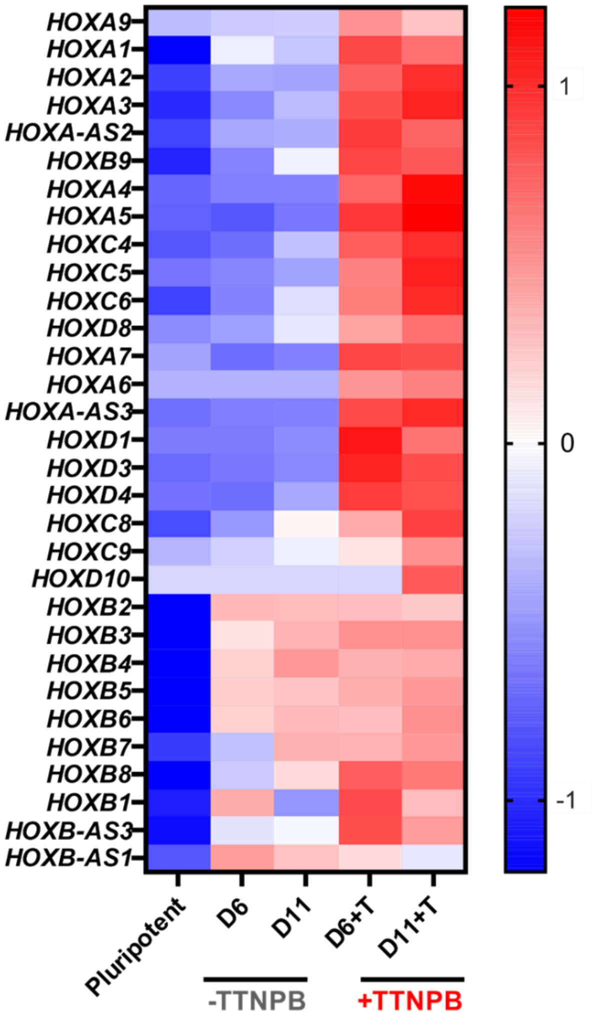


**Supplementary Figure 7 - Analysis of HOX gene expression influenced by TTNPB addition.** Genes centred using clustering analysis software (31). Gene expression was included only if adjusted p value was <0.1 between 1 or more conditions

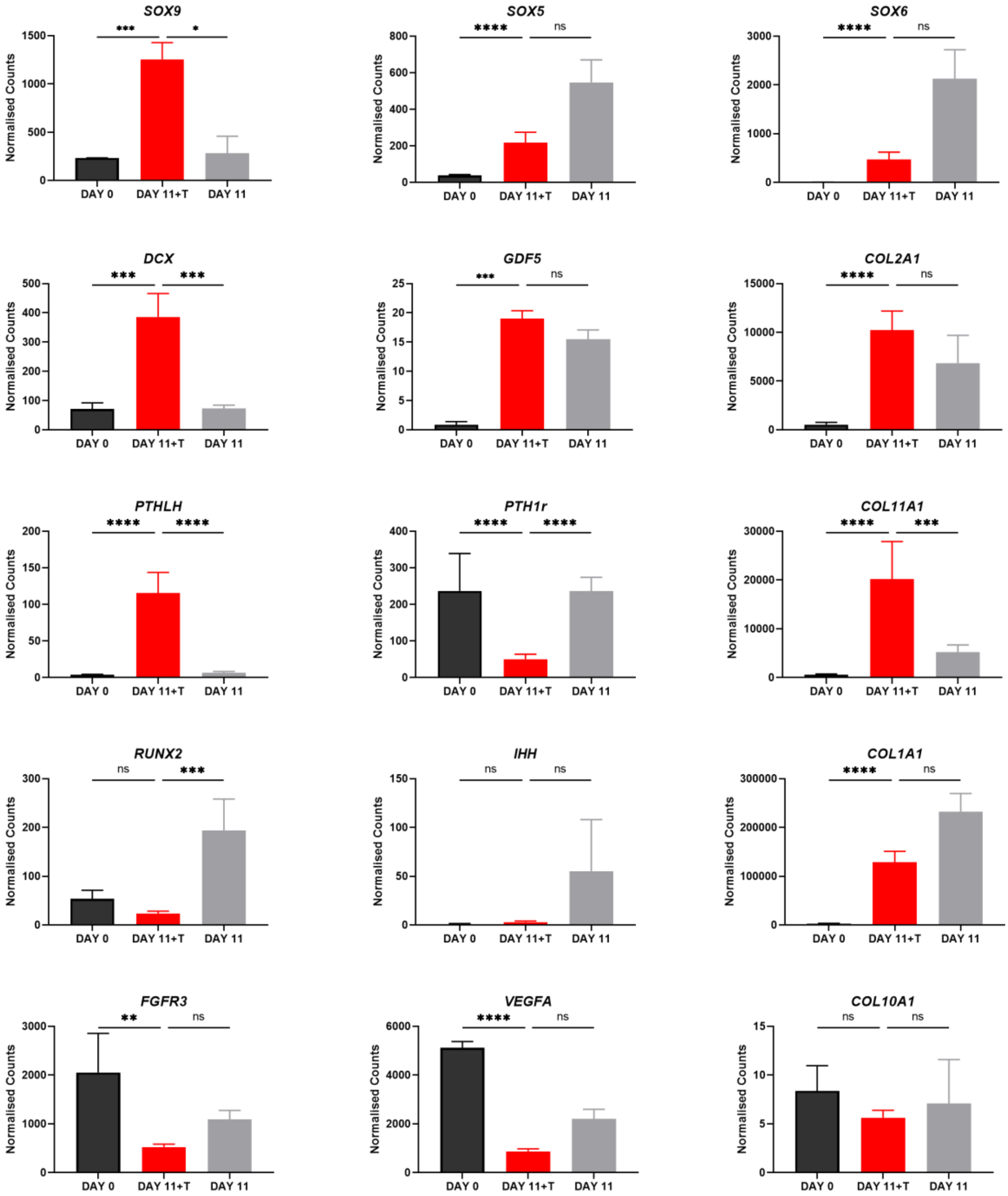


**Supplementary Figure 8 - Gene expression of articular and hypertrophic cartilage markers**. Data shown as MEAN + SEM of N=3 different biological repeats, normalised read counts were taken from the bulk RNA-seq experiment. Statistical differences between DEGs were calculated using DESeq2 and adjusted p values are shown. (*adj. p<0.1, **adj. p<0.05, ***adj. p<0.005, ****adj. p<0.001).

### Supplemental Experimental Procedures

**Supplementary Table 1 – Overview of the RAPID-E differentiation protocol –** For allowing pellet formation, TTNPB supplementation was ceased at day 10.

| STAGE | Mid-primitive streak | Lateral plate mesoderm | Limb Bud Mesoderm | Chondrogenesis | | | 3D Chondrogenesis | Manufacturer |
|---|---|---|---|---|---|---|---|---|
| **DAY** | 1-2 | 3-4 | 5-6 | 7-8 | 9 | 10-15 | 15-28 | |
| **Activin A** | 10 ng/ml | | | | | | | Peprotech, #120-14 |
| **BMP2** | | | | | 50ng/ml | 50ng/ml | 25ng/ml | Peprotech, #120-02 |
| **CHIR** | 3µM | | | | | | | Tocris, #4423 |
| **FGF2** | 20 ng/ml | 10ng/ml | 10ng/ml | 10ng/ml | | | | Peprotech, #100-18B |
| **PIK90** | 100nM | | | | | | | Selleckchem, #S1187 |
| **C59** | | 1µM | | 1µM | | | | Abcam, #ab142216 |
| **SB431542** | | 2µM | | | | | | Tocris, #1614 |
| **GDF5** | | | | 20ng/ml | | 40ng/ml | 20ng/ml | Peprotech, #120-01 |
| **TGFβ3** | | | | | | | 10ng/ml | Peprotech, #100-36E |
| **TTNPB** | 100nM | 100nM | 100nM | 100nM | 100nM | | | Tocris #0761 |
| **SUBSTRATE** | Vitronectin | | Fibronectin | | | | | |
| **MEDIUM** | ADBM | | | | | | ADBM + A2P | |

**Supplementary Table 2 - Overview of antibodies used for immunohistochemistry and the concentration used.**

| Target | Isotype | Dilution | Product code | Secondary antibody |
|---|---|---|---|---|
| Aggrecan | Rabbit Polyclonal | 1:200 | Kind Gift Prof. T Hardingham (92) | Goat anti-rabbit |
| Collagen I | Mouse IgG | 1:500 | Abcam #ab6308 | Horse anti-mouse |
| Collagen II | Mouse IgG | 1:200 | Abcam #ab185430 | Horse anti-mouse |
| Collagen X | Rabbit IgG | 1:200 | Abcam #ab58632 | Goat anti-rabbit |
| Lubricin | Rabbit IgG | 1:200 | Abcam #ab94933 | Goat anti-rabbit |
| SOX9 | Mouse IgG | 1:100 | Abcam #76997 | Horse anti-mouse |

**Supplementary Table 3 – Primers used for RT-qPCR**

| Target Gene | Forward (5' to 3') | Reverse (5' to 3') |
|---|---|---|
| ***GAPDH*** | ATGGGGAAGGTGAAGGTCG | TAAAAGCAGCCCTGGTGACC |
| ***POU5F*** | AGACCATCTGCCGCTTTGAG | GCAAGGGCCGCAGCTTA |
| ***NANOG*** | GGCTCTGTTTTGCTATATCCCCTAA | CATTACGATGCAGCAAATACAAGA |
| ***SOX5*** | ATCCCAACTACCATGGCAGCT | TGCAGTTGGAGTGGGCCTA |
| ***SOX9*** | GACTTCCGCCACGTGGAC | GTTGGGCGGCAGGTACTG |
| ***COL1A1*** | CAGCCGCTTCACCTACAGC | TTTTGTATTCAATCACTGTCTTG |
| ***COL2A1*** | GGCAATAGCAGGTTCACGTACA | CGATAACAGTCTTGCCCCACTT |
| ***COL10A1*** | CAGGCATAAAAGGCCCACTA | AGGACTTCCGTAGCCTGGTT |
| ***ACAN*** | GCAGAGACGCATCTAGAAATTG | GGTAATTGCAGGGAACATCATT |
| ***RUNX2*** | GCCTTCAAGGTGGTAGCCC | CGTTACCCGCCATGACAGTA |
| ***PRG4*** | GAGTACCCAATCAAGGCATTATCA | CCATCTACTGGCTTACCATTGCA |
| ***BMPR1B*** | GAGGATGACTCTGGGTTGCC | AGGCAGTGTAGGGTGTAGGT |
| ***FOXF1*** | AGCAGCCGTATCTGCACCAGAA | CTCCTTTCGGTCACACATGCTG |
| ***HAND1*** | GTGCGTCCTTTAATCCTCTTC | GTGAGAGCAAGCGGAAAAG |
| ***HAND2*** | ACATCGCCTACCTCATGGAC | TTCTTGTCGTTGCTGCTCAC |
| ***ISL1*** | AGATTATATCAGGTTGTACGGGATCA | ACACAGCGGAAACACTCGAT |
| ***NKX-2.5*** | CAAGTGTGCGTCTGCCTTT | CAGCTCTTTCTTTTCGGCTCTA |
| ***PRRX1*** | TGATGCTTTTGTGCGAGAAGA | AGGGAAGCGTTTTTATTGGCT |
| ***HOXB5*** | AACTCCTTCTCGGGGCGTTAT | CATCCCATTGTAATTGTAGCCGT |
| ***TBX6*** | AAGTACCAACCCCGCATACA | TAGGCTGTCACGGAGATGAA |
| ***PAX1*** | CTCACAGCTGGCAGGGTATC | TTAGAGACTGCATGTTAGTTCTGGA |
